# Sphingolipid production by gut Bacteroidetes regulates glucose homeostasis

**DOI:** 10.1101/632877

**Authors:** Elizabeth L. Johnson, Stacey L. Heaver, Jillian L. Waters, Benjamin I. Kim, Alexis Bretin, Andrew L. Goodman, Andrew T. Gewirtz, Tilla S. Worgall, Ruth E. Ley

## Abstract

Levels of Bacteroidetes in the gut microbiome are positively associated with insulin resistance (IR) in humans. Considering that IR is promoted by elevations in hepatic sphingolipids (SL), particularly ceramides, and that Bacteroidetes are the only microbiome phylum possessing genes encoding serine palmitoyltransferase (SPT), which mediates SL synthesis, we investigated a potential link between bacterial SL production, host SL metabolism, and IR. *In vitro*, bacterial SLs entered colonocytes and were metabolized into complex SL, including ceramides. In mice, administration of WT *Bacteroides thetaiotaomicron*, but not a SPT-deficient mutant, resulted in elevated levels of liver ceramides and reduced responsiveness to exogenously administered insulin. This work establishes bacterial SLs as a new class of microbiome-derived molecule capable of impacting host metabolism.

**One Sentence Summary:** SL production by gut Bacteroidetes regulates liver ceramide levels and insulin sensitivity.

## Main Text

Insulin resistance (IR) is a hallmark of metabolic syndrome, the precursor to type 2 diabetes. Ceramides, which are an important class of complex sphingolipids (SLs), have been implicated in the development of IR in animal models (*1*). Reduction of ceramide (d18:1/16:0) in the liver alleviates IR (*2–4*), and pharmacological inhibition of ceramide-related pathways in the intestine leads to improved glucose homeostasis (*5*). In human populations, several studies have associated levels of hepatic or plasma ceramides to IR (*6–9*), suggesting that an understanding of how ceramide pools are regulated could be beneficial for treatment. Sources of SL include *de novo* synthesis, degradation and recycling of existing SLs, and uptake of dietary SLs from the intestine. We hypothesized that the gut microbiome constitutes an additional source of SL with impact on host SL homeostasis.

The human gut microbiome comprises a vast diversity of bacterial species that varies between individuals, however across individuals a small number of bacterial phyla are proportionally dominant. One of the dominant phyla, the Bacteroidetes, has a functionally unique feature that distinguishes them from other members of the gut microbiome: they are able to synthesize SLs, which are very similar, or identical, in structure to mammalian SLs (*10–12*). Members of the phylum Bacteroidetes include genera such as *Bacteroides, Prevotella* and *Porphyromonas*, all of which are highly prevalent in humans, wherein they together constitute 30-40% on average of the gut microbiome (*12*), and therefore possibly act as an endogenous source of SL to their hosts.

Several metagenomic studies have revealed associations between elevated levels of Bacteroidetes in the gut and markers of metabolic disease (*13, 14*). In a study of 292 Danes, Le Chatelier *et al*. noted a reduction of *Bacteroides* in low-richness gut microbiomes associated with hosts with lower insulin resistance (*14*). Pederson *et al*. linked IR and elevated Bacteroidetes in the gut via the production of branched-chain amino acids in 277 non-diabetic individuals (*13*). Bacteroidetes have also been shown to influence IR via the metabolism of bile acids (*15, 16*), and FXR signaling (*5, 17*). Considering that host SLs levels are linked to IR *(6–9)*, and that Bacteroidetes produce SL, these bioactive lipids may provide a mechanistic link between gut Bacteroidetes and IR. Given their large number in the intestine, here we investigate whether Bacteroidetes may be an additional source of SLs to the host with consequences for liver ceramide homeostasis and IR.

To date, evidence that bacterial SLs interact with host tissue comes from the observation that a glucosylceramide produced by *B. fragilis* regulates iNKT cell proliferation in neonatal mice (*18*). To assess the potential for SLs of bacterial origin to interact with non-hematopoietic host cells, we first tested whether bacterial SLs were readily taken up by mammalian cells, and whether bacterial lipids could be processed by mammalian SL synthesis pathways. *Bacteroides spp*. produces both even-chain length sphinganine (Sa - d18:0) and odd-chain length Sa (d17:0)(*10*), while mammalian SPT produces only even-chain Sa (d18:0) (*19, 20*) (for an overview of SL synthesis pathways, see Fig S1). This difference in chain length allows the tracking of the bacterial long-chain base Sa into mammalian cells. We supplemented Caco-2 cells with Sa (d18:0) to stimulate *de novo* SL synthesis (Fig. S2A-B, see methods), then dosed the cells with different amounts of the odd-chain base bacterial Sa (d17:0). We observed the conversion of Sa (d17:0) to So (d17:1), indicating that Sa (d17:0) was readily taken up by Caco-2 cells and incorporated into complex SLs through the *de novo* synthesis pathway (Fig. 1A-B). This observation establishes that odd-chain base SLs of bacterial origin can be processed by the mammalian SL pathways.

**Fig. 1.**
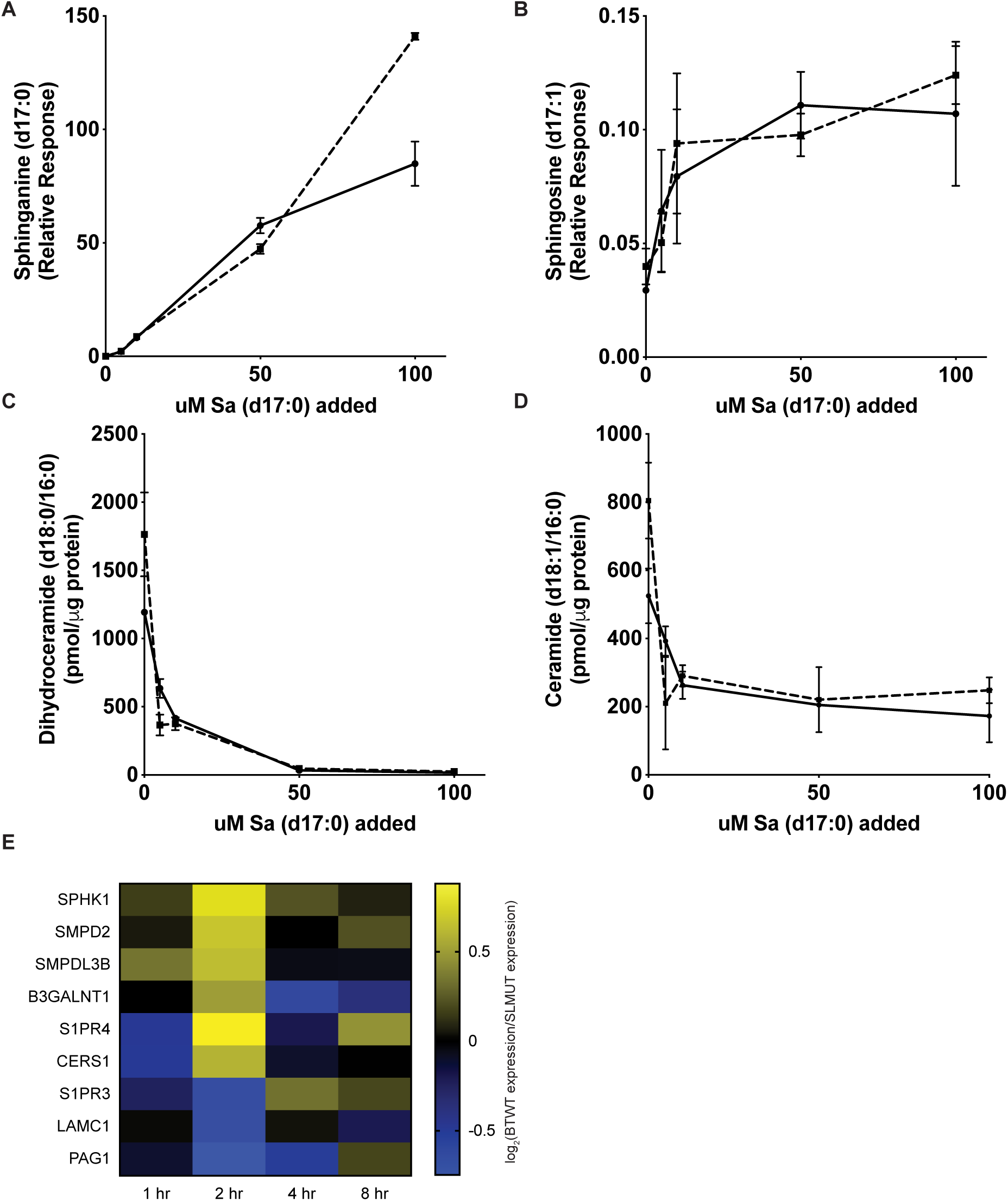
SL production by gut microbes inhibits *de novo* SL synthesis and alters gene expression in Caco-2 cells. (A-D) SL synthesis was induced in proliferating Caco-2 cells with addition of 5 uM Sa (d18:0) and then cells were dosed with increasing concentrations (5, 10, 50, 100 uM) of Sa (d17:0) to monitor the ability of Sa (d17:0) to inhibit flux of C18-base length SLs through the SL synthesis pathway. Cells were harvested 1 hour after addition of lipids and the two curves (dotted and straight lines) represent independent replicates of the same time course. Means ± SEM of LC-MS measurements (n=2) are plotted for (A) Sa (d17:0), (B) So (d17:1), (C) dihydroceramide (d18:0/16:0), (D) ceramide (d18:1/16:0). (E) Heatmap of the average time-zero normalized log_2_ change in gene expression between BTWT and SLMUT in transwell with Caco-2 cells for two replicate 8-hr experiments.

In addition to the processing of bacterial SL by the mammalian pathway, we observed inhibition of Sa (d18:0) metabolism at higher levels of Sa (d17:0). Levels of Sa (d18:0) were lower than baseline at 5µM concentrations of Sa (d17:0), indicating flux of Sa (d18:0) through the SL-processing pathway. However, at higher concentrations of Sa (d17:0), Sa (d18:0) levels remained high (Fig. S2C), which indicates that Sa (d18:0) metabolism stalled even though Sa (d17:0) metabolism was active. Furthermore, we observed a concentration-dependent decrease in the ceramide precursor dihydroceramide (d18:0/16:0 - Fig. 1C) and the downstream products ceramide (d18:1/16:0; Fig. 1D) under increasing Sa (d17:0) conditions, suggesting an overall inhibition of *de novo* C18 long chain base ceramide production. This inhibition of C18 long chain base ceramide production was also observed at lower concentrations of Sa (d17:0) (*i.e*., 0.5-2 µM; Fig. S2E). Levels of sphingomyelin (d18:1/16:0; Fig. S2D) were unchanged with the addition of Sa (d17:0). These results indicate that (i) odd-chain length bacterial Sa (d17:0) can be taken up by mammalian cells, then (ii) metabolized by SL-processing pathways *in vitro*, while (iii) inhibiting the production of mammalian C18-base SLs.

We next assessed if lipids could transfer from bacterial cells to mammalian cells. We used *Bacteroides thetaiotaomicron* VPI 5482 (hereafter, BTWT) cells grown in medium supplemented with palmitic acid alkyne (PAA; allowing for fluorescence tracking using click chemistry) in a transwell system, wherein bacterial cells were located 1 mm above mammalian cells and separated by a bacterial-impermeant 0.4 uM pore. Using this system, we observed that alkyne-labeled lipids from BTWT were transferred to Caco-2 cells within 4 hours (Fig. S4).

To assess if SL production by BTWT could influence SL metabolism in Caco-2 cells, we exposed them to BTWT and the same strain unable to synthesize SLs due to an inactivated SPT gene (hereafter, SLMUT; Fig. S3; Methods) (*21, 22*). We performed a transcriptome analysis of Caco-2 cells exposed to BTWT or SLMUT over 8 hours in the transwell (targeted analysis of 107 SL-related genes, Fig 1E, Fig. S5, Table S1). As expected, genes involved in SL metabolism were differentially regulated in the presence of BTWT or SLMUT. For instance, the sphingosine-1-phosphate receptor gene S1PR4 was upregulated early in Caco-2 cells co-incubated with BTWT compared to SLMUT. S1PR4 serves as a receptor for gut microbiome produced N-acyl amides, and has been suggested to respond to microbiome derived sphingolipids (*23*). Additional genes encoding enzymes in the SL-processing pathway, and upregulated by Caco-2 cells in response to BTWT compared to SLMUT, included a sphingosine kinase (SPHK1), sphingomyelinases (SMPD2, SMPDL3B), and a ceramide synthase (CERS1). SPHK1 is involved in the phosphorylation of the long chain base sphingosine to sphingosine-1-phosphate, and SMPD2 and SMPDL3B in the hydrolysis of the phospholipid sphingomyelin into ceramide and phosphocholine (*24*). CERS1 produces ceramides with a C18:0 acyl chain length shown to influence obesity-related IR in skeletal muscle (*25*). The differential regulation of SL-metabolism genes is consistent with the different lipid exposure from BTWT and SLMUT.

To assess if lipid transfer from bacterial cells to host cells also occurs *in vivo*, BTWT-PAA cells were administered by oral gavage to 5-week old germfree Swiss-Webster (SW) mice. PAA-metabolites, readily observed in bacterial cells, transferred from the lumen into host gut epithelial tissue (Fig. 2A-D). We then introduced BTWT cells containing Sa (d17:0) and Sa (d18:0) in equivalent amounts (Fig. S6) via daily oral gavage to germfree SW mice over a one-week period and measured SLs in host tissue over the course of the week. On days 1 and 7, we measured higher Sa (d17:0) in hepatic portal vein blood in BTWT-treated compared to untreated mice (Fig. 2E). Together these results establish that *Bacteroides*-SLs can transit from bacterial cells to the gut epithelium and portal vein, which should allow them to reach the liver.

**Fig. 2.**
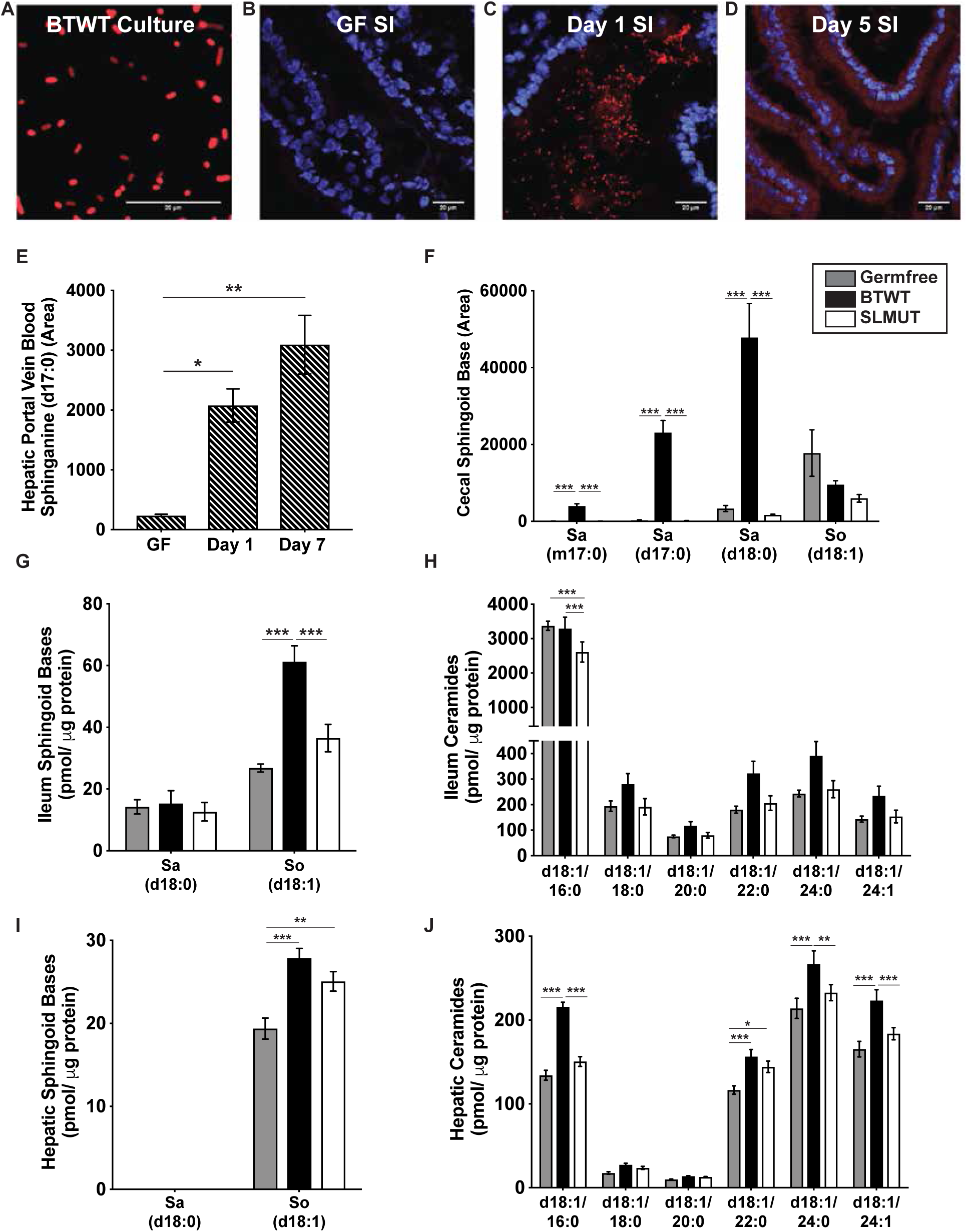
SL production by BTWT results in intestinal lipid uptake, increased odd chain SLs in the hepatic portal vein circulation, and increases in cecal, ileal, and hepatic SL levels. Confocal microscopy of (A) BTWT cells grown in minimal media supplemented with palmitic acid alkyne (PAA), (B) small intestine (SI) tissue of germfree mice, (C) SI tissue of germfree mice inoculated with BTWT tagged with PAA 3 hours after oral gavage, and (D) SI tissue of germfree mice after 5 days of daily gavage with BTWT tagged with PAA. PAA metabolites were labeled with Alexa Fluor 647 azide (red) using click chemistry, and nuclei of the intestinal epithelial cells were labeled with DAPI (blue). Scale bar is 20 µm. Representative images of 4 mice. (E) Sphinganine (d17:0) levels in acid base-treated hepatic portal vein blood of germfree mice gavaged daily with BTWT. For hepatic portal vein blood samples, means ± SEM of LC-MS measurements are plotted for: GF=germfree (n=2), one day of daily gavage, Day 1 (n=3); 7 days of daily gavage, Day 7 (n=3) (one-way ANOVA, Tukey’s multiple comparison test, *=p<0.05, **=p<0.01). Long chain base SLs in the (F) cecal content of germfree mice (grey), germfree mice monoassociated with BTWT (black), and germfree mice monoassociated with SLMUT (white). Sphingoid base (G) and ceramide (H) levels in ileum tissue of germfree mice (grey), germfree mice monoassociated with BTWT (black), and germfree mice monoassociated with SLMUT (white). Sphingoid base (I) and ceramide (J) levels in hepatic tissue of germfree mice (grey), germfree mice monoassociated with BTWT (black), and germfree mice monoassociated with SLMUT (white). (F – J) Bar charts represent mean SL abundance ± SEM for 12 mice per condition (GF, SLMUT) and 11 mice per condition (BTWT), (two-way ANOVA, Tukey’s multiple comparison test, *=p<0.05, **=p<0.01, ***=p<0.001).

Conventionalization of germfree mice results in a normalization of lipid cycling, including SL metabolism (*26*). Previous reports have shown that (i) germfree mice have higher hepatic ceramides compared to conventionally-raised mice, and that (ii) conventionalization of germfree mice by inoculation with a complex microbiota normalizes hepatic ceramides (*27, 28*). We confirmed these results by conventionalizing GF mice and comparing liver ceramides to those of conventionally-raised animals (Fig. S7). Moreover, to dissect how host hepatic ceramide levels relate specifically to the SL-production capacity of the gut microbiota, we colonized 4 week old germfree SW mice with either BTWT or SLMUT for 6 weeks and profiled SLs in tissues. BTWT and SLMUT colonized the mouse cecum at comparable levels (Fig. S8A). At 6 weeks post colonization, cecal levels of three long chain base SLs synthesized by BTWT, Sa (d17:0), 15-methylhexadeca Sa (m17:0), and Sa (d18:0), were significantly higher in BTWT-colonized mice compared to SLMUT-colonized and GF mice, whereas levels of the long chain base SL So (d18:1), which is not synthesized by BTWT, did not differ between treatments (Fig. 2F). Cecal dihydroceramides and ceramides with longer acyl chains (C22:0, C24:0, C24:1) were elevated in the SLMUT condition, but cecal sphingomyelins were similar between conditions (Fig. S8B-D). Thus, the SL-production capacity of the microbes in the gut has a large influence on the SL-milieu of the lumen.

Consistent with uptake and processing of saturated SL bases (d18:0) from the lumen, ileal levels of the long chain base So (d18:1) were higher in BTWT compared to SLMUT and GF conditions (Fig. 2G). Dihydroceramide (d18:0/18:1), and dihydroceramide (d18:0/24:0) (Fig. S8E) were also significantly elevated in the BTWT condition compared to SLMUT and GF conditions. Additionally, ileal ceramide levels (d18:1/18:0, d18:1/20:0, d18:1/22:0, d18:1/24:0, d18:1/24:1) trended higher in the BTWT compared to SLMUT and GF conditions (Fig. 2H), while ceramide and sphingomyelin (d18:1/16:0) were higher in the GF and BTWT conditions compared to the SLMUT (Fig. S8F). These observations suggest that saturated SL long chain bases (Sa (d18:0) and Sa (d17:0)) are taken up from the lumen by intestinal epithelial cells and metabolized through the host’s *de novo* synthesis pathway in the intestine, resulting in increased abundance of the downstream products sphingosine, dihydroceramide, and ceramide.

Importantly, hepatic dihydroceramide and ceramide levels were significantly higher in the BTWT compared to SLMUT for multiple prominent ceramide species. Specifically, hepatic levels of dihydroceramide (d18:0/24:1) (Fig. S8G), ceramide (d18:1/16:0), ceramide (d18:1/24:0), ceramide (d18:1/24:1) (Fig. 2J), sphingomyelin (d18:1/16:0), and sphingomyelin (d18:1/24:1) were individually higher in BTWT compared to SLMUT and GF conditions (Fig. S8H). These results indicate that bacterial production of SL in the gut can impact levels of liver ceramides.

We next tested whether oral administration of BTWT to conventionally-raised mice could supplement a diet deficient in SLs and impact rates of *de-novo* synthesis. We placed 5-week old female SW mice on a fatty-acid free (FAF) diet known to increase hepatic lipogenesis (*29*). After 3 weeks on the diet, hepatic *de novo* SL synthesis (as measured by the increase in the ratio of hepatic dihydroceramides compared to ceramides) was elevated in the livers of FAF-fed mice compared to controls on breeder chow (Fig.S9A). Mice were gavaged twice with BTWT or SLMUT over two days. BTWT treatment was associated with the highest hepatic ceramide levels (Fig. 3A). Of the ceramides detected in the liver, the most abundant was (d18:1/24:1) (Fig S9B). We observed significantly reduced *de novo* synthesis of this abundant ceramide in the BTWT compared to the SLMUT and PBS treatments (Fig. 3B). The SLMUT-gavage yielded effects on *de novo* SL synthesis similar to those of the PBS-gavage control (Fig. 3B). Thus, when mice are fed a SL-poor diet, gut bacteria can act as an endogenous source of SLs that reduces *de-novo* synthesis.

**Fig. 3.**
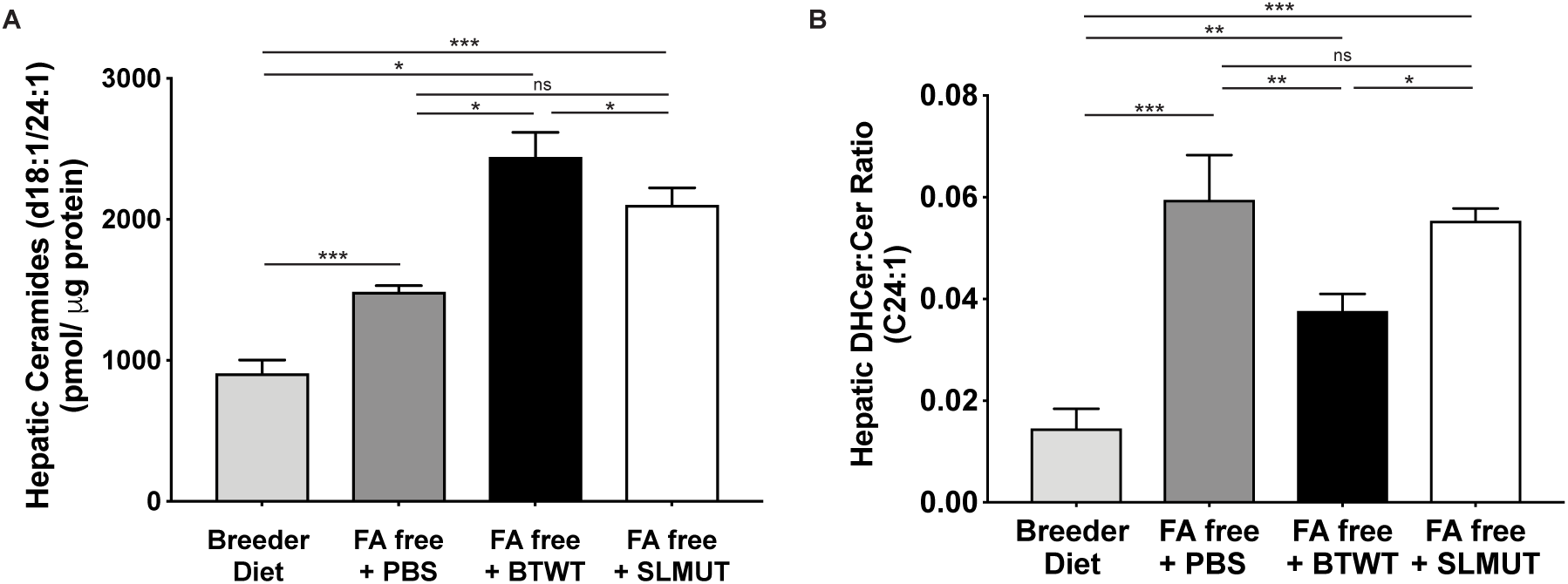
Oral supplementation of BTWT results in increased hepatic ceramides and inhibition of hepatic *de novo* SL synthesis *in vivo*. (A) Total hepatic ceramides in mice on a breeder diet or a fatty acid free (FA free) diet that stimulates hepatic *de novo* SL synthesis. Mice on the FA free diet were gavaged with a PBS control, BTWT, or SLMUT. (B) Hepatic dihydroceramide (DHCer) to ceramide (Cer) ratios for Ceramide (d18:1/C24:1). Mean values ± SEM from one experiment are plotted, n=7 per condition, (one-way ANOVA, Tukey’s multiple comparison test, *=p <0.05, ***=p<0.001, ns = not-significant).

Elevated hepatic ceramide levels have the potential to affect insulin sensitivity in the host (*2–4, 29, 30*). We assessed how dietary supplementation with BTWT affects hepatic ceramides in high-fat diet fed C57BL/6J mice, a well-established mouse model of IR. Conventionally-raised C57BL/6J mice originally on a chow diet were gavaged with BTWT or SLMUT (in this instance, an SPT knock-out strain, Methods) for 7 days, switched to a high-fat diet and re-gavaged with the same bacterial treatment for 7 days, then maintained on the high-fat diet for 21 days. Mice were then switched to back to a chow diet and gavaged with the same strains daily for 9 days. Relative to the SLMUT-treated mice, the BTWT-treated mice exhibited higher hepatic levels of ceramide (d18:1/16:0) and ceramide (d18:1/18:0) (Fig. 4A), sphinganine (d18:0), and sphingosine (d18:1) (Fig. 4B). Moreover, in an insulin tolerance test, the BTWT-treated mice exhibited lower insulin sensitivity compared to SLMUT-treated mice (Fig. 4C), although a glucose tolerance test indicated a similar glucose handling response (Fig. 4D). Our observed effects of BTWT dosing on IR are similar to what has been observed when mice are treated with other members of the Bacteroidetes, such as *P. copri* (*13*) and *B. fragilis* (*17*). Our results further indicate that the BTWT treatment resulted in elevated hepatic ceramides, with the expected associated reduced insulin sensitivity.

**Fig. 4.**
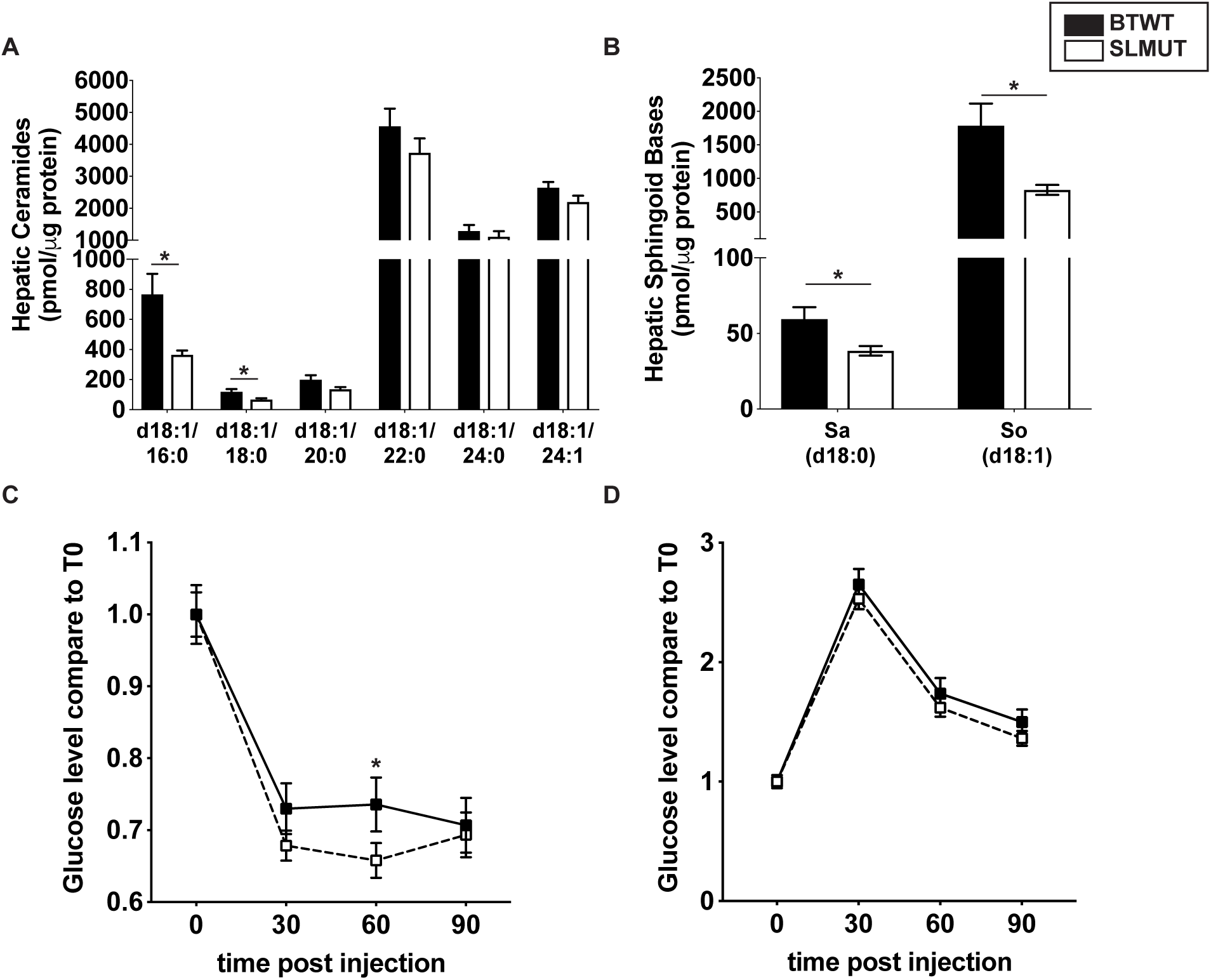
Increased hepatic ceramides and insulin resistance in mice supplemented with BTWT as compared to SLMUT. Mice fed normal chow were gavaged with BTWT or SLMUT and then switched to a high fat diet and gavaged with BTWT or SLMUT. Mice were then kept on a high-fat diet for 21 additional days to induce insulin resistance and switched to back to normal chow. These mice were then gavaged with BTWT or SLMUT for 9 days before SL assessment and insulin phenotyping. Insulin resistance was induced in mice and SL levels and insulin phenotypes were assessed. Hepatic ceramides (A), long chain base SLs (B), insulin tolerance (C), and glucose tolerance (D) were measured at the conclusion of the experiment. (A-D) Mean values ± SEM are plotted, n=10 per treatment, combined data from two replicate experiments with similar trends. (A-B) Observations with significant differences in SL abundance or glucose levels between treatments are marked with stars (t-test, *=p <0.05). (C) T-test for glucose levels at 30-min, (*=p <0.05).

This study points to a role for SL production by Bacteroidetes to modulate the levels of bioactive lipids in the liver and in turn affect IR. We have demonstrated that mammalian cells sense and respond to the presence of bacterial SLs, and that these SLs can be processed via mammalian SL pathways. Furthermore, our results show that mice adjust hepatic *de-novo* production of sphingolipids in response to the SL-production capacity of gut bacteria. These observations provide an additional mechanistic link that may underlie the observed positive associations between the levels of Bacteroidetes in the gut and IR. Given that all members of the Bacteroidetes phylum produce SLs, and that the Bacteroidetes constitute a widespread and dominant phylum in the human gut, levels of these bacteria may eventually be factor to consider in the management of IR.

## Acknowledgements

We thank Noah J. Clark, Qiaojuan Shi, Hylde Zipoli, Liz Chang, Inge Hansen and Richard Deckelbaum.

## Author Contributions

E.L.J, T.S.W, R.E.L conceived of the experiments. E.L.J, S.L.H, J.K.G, A.L.G, T.S.W, R.E.L designed the experiments. E.L.J, S.L.H, B.I.K, A.B, and J.L.W performed the experiments. E.L.J analyzed the data. E.L.J and R.E.L. wrote the manuscript with help from S.L.H and T.S.W.

## Data and materials availability

All processed data is available in the main text or supplementary materials.

## Supplemental Materials and Methods

### Caco-2 cell culture

Human epithelial colorectal adenocarcinoma cells (Caco-2, ATCC) were cultured in Dulbecco’s modified Eagle’s medium (DMEM, ThermoFisher Scientific) supplemented with 10% fetal bovine serum (FBS, Gibco). Plates were seeded at 5 × 10^5^ cells per 10 cm plate, incubated at 37°C with 5% CO_2_, and experiments took place 48 hours after the initial seeding. For lipid competition experiments, sphinganine (d18:0) (Avanti Polar Lipids) was added to the media at a concentration of 5 µM. Sphinganine (d17:0) (Avanti Polar Lipids) was also added to plates at concentrations of 0 µM, 5 µM, 10 µM, 50 µM, and 100 µM. The lipid competition experiment was done in duplicate and repeated a third time using a lower stimulating concentration of sphinganine (d18:0) of 1 µM with the addition of sphinganine (d17:0) at concentrations of 0.5 µM, 1 µM, and 2 µM. Plates in which there was a media change but no addition of lipid were also collected. Cell pellets from plates were collected after a 1-hour incubation and stored at - 80°C.

### Bacterial culturing

*Bacteroides thetaiotaomicron* strain VPI 5482 (wild type, BTWT) and the SLMUT strains are described in (*21*). SLMUT consists of BTWT with a transposon insertion in the gene BT0870, which is annotated to have 8-amino-7-oxononanoate synthase activity and is homologous to a gene with serine palmitoyl transferase (SPT) activity in *Bacteroides fragilis (22)* and the known SPT in yeast. The insertion is in a position 88% from the start of the gene. Unless otherwise stated, BTWT and SLMUT strains used in experiments described below were grown under anaerobic conditions at 37°C in either chopped meat broth (ThermoFisher Scientific/Remel), supplemented brain heart infusion media (BHIS), or a minimal medium with glucose as the sole carbon source (MMG) (*31*). The minimal medium consisted of, per L: 13.6 g KH_2_PO_4_, 0.875g NaCl, 1.125 g (NH_4_)_2_SO_4_, 5 g glucose, (pH to 7.2 with concentrated NaOH), 1 mL hemin solution (500 mg dissolved in 10 mL of 1M NaOH then diluted to final volume of 500 mL with water), 1 mL MgCl_2_ (0.1M in water), 1 mL FeSO_4_x7H_2_O (1 mg per 10 mL of water), 1 mL vitamin K3 (1 mg/mL in absolute ethanol), 1 mL CaCl_2_ (0.8% w/v), 250 µL vitamin B12 solution (0.02 mg/mL).

### Generation of BT0870 knockout strain (Δ0870-SLMUT)

To generate an in-frame deletion of BT0870 in *B. thetaiotaomicron*, the strain *B. thetaiotaomicron* VPI-5482 *tdk* was used as previously described (*32*). Two 800-1000 bp regions, each flanking the gene to be deleted, were PCR-amplified (NEB Q5 Hot Start High-Fidelity DNA Polymerase) and cloned into EcoRV-HF and NotI-HF linearized pExchange_tdk. Strains and primers are listed in Table S3. The assembled construct was transformed into *E. coli* S17-1 λpir (Biomedal), plated on LB agar-streptomycin-carbenicillin plates, and transformants screened for incorporation of the plasmid. Final concentrations of antibiotics and selection agents were as follows: erythromycin 25 μg/ml, gentamicin 200 μg/ml, streptomycin 100 μg/ml, carbenicillin 100 μg/ml, FUdR 200 μg/ml. To conjugate, recipient and donor cells were inoculated from overnight cultures (*B. thetaiotaomicron tdk* at 1:1000; E. coli transformant at 1:250) and grown to early exponential phase (OD600 0.2-0.3), upon which the donor and recipient strains were combined in a 1:1 ratio and centrifuged for 20 min at 4000 RPM at room temperature. The bacterial pellet was resuspended in 100 μl BHIS, plated as a puddle on BHIS-10% defibrinated sheep blood agar plates, and incubated aerobically at 37°C for 20 hours. The conjugation puddle was then scraped, serially diluted in PBS, and incubated aerobically at 37°C on BHIS-10% defibrinated sheep blood-gentamicin-erythromycin agar plates. Colonies were screened for merodiploids via PCR, cultured overnight in liquid BHIS, and serially diluted onto BHIS-10% defibrinated sheep blood-gentamicin-FUdR agar plates. Colonies were PCR screened for deletion of the gene and confirmed via Sanger sequencing. This strain is differentiated from the transposon insertion mutant (SLMUT) as Δ0870-SLMUT and was used in the high-fat diet feeding experiments (below).

### Thin Layer Chromatography (TLC)

To resolve sphingolipid species, lipid extracts were spotted on non-fluorescent 20 × 20 cm glass backed silica plates (Millipore Sigma) and separated using a 65:25:4 Chloroform: Methanol: Ammonium Hydroxide solvent system for 25 minutes. A standard composed of sphingosine (d18:1), ceramide (d18:1/12:0), and sphingomyelin (d18:1/12:0) (Avanti Polar Lipids) was used to identify spots. All plates were developed in an iodine chamber overnight.

### Inhibition of sphingolipid synthesis in BTWT

BTWT and SLMUT were grown in BHIS supplemented with 1µM of myriocin (Sigma-Aldrich), an inhibitor of SPT, to inhibit *de novo* sphingolipid synthesis. Lack of sphingolipid species in SLMUT was confirmed by both liquid chromatography-mass spectrometry (LC-MS) and TLC analysis of lipid extracts (Fig. S2A-C).

### Outer membrane vesicle (OMV) preparation

BTWT and SLMUT cultures were separated into fractions (whole cell, cell membranes, outer membrane vehicles (OMVs)) and sphingolipid composition was evaluated by LC-MS. To separate the cell fraction (for whole cells and membranes) from the supernatant fraction (for OMVs), 100 mL of 18-hr bacterial cultures grown at 37°C in minimal media were spun at 3220 x g for 20 min, the supernatant was moved to a clean tube, and the spin repeated. To prepare OMVs, the supernatant was filtered twice using a 0.22 µm pore membrane (Corning Inc.). 60 mL of each culture was spun at 140,000 x g for 2 hours at 4°C. OMV pellets were washed in PBS and spun at 140,000 x g for 2 hours at 4°C, then resuspended in 100 µL PBS and stored at −80°C prior to lipid extraction. Purity of the OMV fraction was confirmed by negatively staining the OMV preparation and imaging by transmission electron microscopy at the Cornell Center for Materials Research at Cornell University. For membrane extraction, half of each bacterial pellet was resuspended in 10 mL membrane extraction buffer (MEB; 50 mM Tris-HCl, 150 mM NaCl, 50 mM MgCl_2_, pH 8.0). The suspension was sonicated in 2 × 30 second intervals, then spun 500 x g for 10 minutes at 4°C. The supernatant was spun at 140,000 x g for 2 hours at 4°C, then the membrane fraction pellet was washed in PBS and respun at 140,000 x g for 2 hours at 4°C. Pellets were resuspended in 1 mL PBS and stored at −80°C prior to lipid extraction (*33*). Lipids were extracted from each fraction using the method of Bligh and Dyer (*34*) and acid-base treated as described below (“Measurement of total sphingolipid levels using acid base hydrolysis”). Lipid films were resuspended in 1:1 dichloromethane:methanol prior to analysis by LC-MS.

### Caco-2 transwell incubation with Bacteroides thetaiotaomicron strains

To assess the effects of metabolite transfer from BTWT to intestinal epithelial cells, Caco-2 cells were incubated with BTWT and SLMUT in tissue culture dishes (Corning Inc.), wherein bacterial cells were placed on a 0.4 µM filter situated 1 mm above the monolayer of Caco-2 cells. 50 mL cultures (OD_600_ of 0.25) grown in minimal medium supplemented with 25 µM palmitic acid alkyne (PAA, Cayman Chemical) were washed 3 times in PBS and resuspended in 6 mL of 10% FBS in DMEM. 1 mL of bacterial suspension was added to upper-well inserts of the 6-well transwell culture dished. Contents of the upper bacterial insert and the lower Caco-2 cell filled plate were collected 4 hours after the addition of the bacteria. Alkyne-containing metabolites in bacterial and Caco-2 cells (plated on UV-sterilized #1.5 glass coverslips in 6-well plates) were labeled with Alexa Fluor 647 using the Click iT cell reaction buffer kit (Thermo Fisher Scientific). Bacterial cells were mounted onto glass slides and imaged on a LSM 710 confocal microscope (Zeiss).

### RNA-seq of Caco-2 cells in transwell with Bacteroides thetaiotaomicron strains throughout an 8-hour time course

RNA was isolated from Caco-2 cells incubated in transwell plates with BTWT or SLMUT cultures using the Trizol reagent (ThermoFisher) according to manufacturer’s instructions. Samples were collected from BTWT and SLMUT conditions over an 8-hour time period (0, 1, 2, 4, 8 hours) and the time course was performed twice. The NEB Next Ultra RNA library kit for Illumina (NEB) was used to prepare sequencing libraries, which were sequenced using 50 bp single end reads on the Illumina HiSeq 2500 platform. The 18 libraries (duplicates of BTWT and SLMUT 1, 2, 4, and 8-hour timepoints (16 libraries) in addition to duplicates of the time 0 timepoint (2 libraries)) were multiplexed using Illumina barcodes and read across two lanes of a flow cell (9 libraries per lane) for a total of 474,824,584 reads and an average of 26,379,143 reads per sample. Reads were mapped to the human genome using the STAR aligner (*35*) with an average of 85% (range 80 – 87%) of total sequenced reads mapping to the transcriptome. Whole transcriptome analysis of differential expression patterns over the time course was done using the maSigPro package implemented in R (*36*). Analysis of the expression of sphingolipid processing genes was done with a set of 107 manually curated genes from gene ontology categories involved in sphingolipid, ceramide, sphingosine, or sphingomyelin metabolism and are detailed in Table S3. For heatmaps, expression values were averaged across replicates and genes with a greater than 2-fold change in normalized expression values are visualized in Fig. 2D.

### Animal experiments

All experiments involving animals were performed according to Protocol #2010-0065 approved by the Cornell University Institutional Animal Care and Use Committee. All gavages of bacterial cultures were prepared from overnight cultures washed and resuspended in sterile PBS and administered in a volume of 0.2 mL at a concentration of 10^8^ CFU/mL unless otherwise noted.

### Daily gavage of BTWT into mice to visualize transfer of fluorescently labeled bacterial lipids to mouse epithelial cells

5-6-week old female germfree Swiss Webster (GF SW) (Taconic Biosciences) were purchased and shipped to Cornell University, where they were allowed to acclimate for 48 hours in their shipper. Upon removal from the shipper, mice were either immediately sacrificed or gavaged with overnight cultures of BTWT grown in MMG supplemented with 25 µM PAA. Mice were then housed 3 – 4 mice per cage and fed an autoclaved breeder diet (5021 LabDiet) *ad-libitum*. Mice were gavaged daily with the same dose of PAA-labeled BTWT for 4 additional days until sacrifice. On the day of sacrifice, mice were fasted for 6 hours then gavaged with PAA-labeled BTWT (as above) 1 hour before sacrifice. Intestinal tissue was harvested for confocal imaging as described below.

#### Bacterial cells

Alkyne containing metabolites in bacterial cells (plated on UV-sterilized #1.5 glass coverslips in 6-well plates) were labeled with Alexa Fluor 647 using the Click iT cell reaction buffer kit (Thermo Fisher Scientific). Bacterial cells were mounted onto glass slides and imaged on an LSM 710 confocal microscope (Zeiss).

#### Animal tissue

At the time of sacrifice, small intestinal tissue was cut into three equal sections. Then, 4 cm of tissue from the duodenum, jejunum, ileum, in addition to the whole length of the colon were embedded in O.C.T media and snap-frozen using isopentane and liquid N_2_. Blocks were cryosectioned into 8 µm sections, set and fixed on a glass slide using 4% paraformaldehyde. Slides were stained using the Click iT cell reaction buffer kit (ThermoFisher Scientific) and imaged on an LSM 710 confocal microscope (Zeiss) at the Cornell University Biotechnology Resource Center.

### Daily gavage of BTWT to GF mice to measure hepatic portal vein blood uptake of sphingolipids

To assess intestinal sphingolipid uptake, 5-week GF SW mice (Taconic Biosciences) were allowed to acclimate for 2 days in their germfree shipping container. After this acclimation period, mice were either immediately sacrificed or gavaged with BTWT cultures grown overnight in MMG. Mice were housed in sterile filter top plastic cages under specific pathogen free (SPF) conditions and sacrificed 6 hours post-gavage on day 1, and 7 of the experiment. Blood was collected from the hepatic portal vein by euthanizing the mice using CO_2_ followed by cervical dislocation. Heparin (100 uL - 30 IU/mL, Sigma) was added to the cavity before nicking the hepatic portal vein and blood was collected using a Pasteur pipet. Hepatic portal vein blood was frozen in liquid N_2_ and stored at −80°C.

### Monoassociation of GF mice with BTWT or SLMUT

GF SW mice were bred in-house and caged in rigid sterile isolators. Mice (females, 3-4 weeks old) were transferred to flexible bubble isolators and inoculated with either BTWT or SLMUT by oral gavage. Mice were housed 3 – 4 per cage, fed an autoclaved breeder diet (5021, LabDiet) *ad-libitum* and sterility was checked biweekly. Mice were sacrificed 6 weeks after inoculation. After decapitation, livers, PBS flushed ileum tissue, and cecal contents were collected, flash frozen in liquid N_2_ and stored at −80°C until processed for sphingolipid analysis. Colonization efficiency was monitored by determining the colony forming units per gram (CFU/g) of cecal content of sacrificed mice. 50 mg of cecal content per sample (7 samples per condition) was weighted and added to 1 mL of BHIS. Serial dilutions (1:10) of the slurry were made to the dilution 1:10^8^. Serial dilutions were plated on BHIS agar plates and incubated at 37°C in an anaerobic chamber overnight before counting colonies.

### Germfree, conventionalized, and conventionally-raised mice used in hepatic sphingolipid profiling

#### Germfree

GF SW mice (female) were bred in-house and caged in rigid sterile isolators. GF mice were sacrificed at 5 weeks of age.

#### Conventionalized mice

5-week old female GF SW mice were inoculated with 0.2 mL of fecal slurry made from 3 mouse pellets homogenized in 3 mL of sterile PBS. Pellets were obtained from SPF Swiss Webster mice housed in the conventional corridor of the Cornell Mouse Facility. Conventionalized mice were housed in sterile filter top cages and sacrificed a week after colonization.

#### Conventionally raised

5-week old conventionally-raised female mice were obtained from litters two generations after breeding pairs from the germfree colony had been conventionalized.

For all mice: after euthanasia by decapitation, livers were collected, flash frozen in liquid N_2_ and stored at −80°C until processed for sphingolipid analysis.

### Induction of hepatic *de novo* SL synthesis using a fatty acid free diet with short term supplementation of BTWT or SLMUT in mice

5-week old female SW mice (Taconic Biosciences) were fed either a breeder (5021 LabDiet: calories from protein - 23%, carbohydrates - 53%, fat - 24%) or a fatty acid free diet (TD.03314 Teklad: calories from protein - 24%, carbohydrates - 76%, fat - 0%) *ad-libitum* for 21 days while housed in SPF conditions with 2 to 5 mice per cage. Mice were gavaged with BTWT or SLMUT 24 hours before sacrifice, and then again prior to fasting (6 hours before sacrifice). After decapitation, livers were collected, flash frozen in liquid N_2_ and stored at −80°C until processed for sphingolipid analysis.

### Insulin-resistant mice administered BTWT or SLMUT

7-week old male C57BL/6J mice (Jackson Laboratory) on a chow diet (Diet 5001, LabDiet) were gavaged with BTWT or SLMUT for 7 days and then placed on a high-fat diet (HFD; D12492, Research diet, Inc.). Mice were then gavaged for 7 additional days with BTWT and SLMUT and kept on the HFD for an additional 21 days. When mice were then switched back to a chow diet and gavaged daily with BTWT or SLMUT for 9 days before sacrifice. All gavages were 0.2 mL of cultures of BTWT or Δ0870-SLMUT (10^8^ CFU/mL) grown in BHIS and washed in PBS. Glucose and insulin tolerance were measured weekly and directly before sacrifice. Briefly, mice were fasted for 5h and fasting glucose level were determined using a Nova Max plus Glucose meter. Then mice were injected with 2mg of glucose/gm of body weight or 0.5 U insulin/kg of body weight and blood glucose levels were measured at 30, 60, 90 min after injection. Mice were euthanized by cervical dislocation and livers were frozen in liquid N_2_ until processed for sphingolipid analysis.

### Lipid extractions

#### Liver, Ileum and Colon sample preparations

Liver, ileum, and colon tissue were all homogenized in PBS using tubes with sterile 1 mm zirconium beads (OPS diagnostics) in a bead beater homogenizer (BioSpec products) for 2 minutes. Intestinal tissue was placed on ice and then subjected to another round of bead beating to ensure homogeneity of the sample. Protein concentrations of homogenates were measured using a Lowry protein assay (BioRad) and equal concentrations of samples were loaded on to 96-well plates for lipid extractions. Liver samples were extracted in 1:1 dichoromethane:methanol according to the details below with 400 – 800 µg of protein per sample while colon and ileum samples were extracted with 100 – 200 µg of protein per sample.

#### Hepatic Portal Vein Blood sample preparations

Hepatic portal vein whole blood (150 µL) was loaded on to 96-well plates for lipid extraction.

#### Cecal tissue preparations

Cecal tissues were weighed before lipids were extracted according to the Folch method (*37*) in 4 mL of 2:1 chloroform:methanol. After 2 hours of constant vortexing on a plate vortexer, 800 µL of 0.9% sodium chloride in water solution was added to the extraction. Samples were briefly vortexed again before spinning samples at 2000 x g for 15 minutes to separate the aqueous and organic phases. The lower organic phase was then transferred to a new tube where 800 µL of 0.9 % sodium chloride was added in order to ensure an organic phase free of cecal debris. These extractions were spun again at 2000 x g for 15 minutes and the organic phase was transferred to a glass tube and dried under nitrogen gas. The remaining lipids were weighed and resuspended at an equal concentration for subsequent mass spectrometry analysis.

#### Cell culture sample preparations

Cell pellets were resuspended in 100 µL of PBS and protein concentrations were assessed using the Lowry method. Equal amounts of cell suspension as measured by protein concentration (100 – 150 µg) were loaded onto 96 well plates for sphingolipid extractions.

#### Bacterial culture sample preparations

Bacterial cell pellets were washed and resuspended in PBS and protein concentrations were assessed using the Lowry method. Equal amounts of cell suspension as measured by protein concentration were loaded onto 96 well plates for sphingolipid extractions.

#### Sphingolipid extraction

All samples loaded onto 96-well plates had 50 µL of 1 µM C12 ceramide (d18:1/12:0) (Avanti Polar Lipids) added as an internal standard and 50 µL of 10% diethylamide diluted in methanol. For the lipid extraction, 450 µL of 1:1 dichloromethane:methanol was added to each sample and vortexed overnight on a plate shaker. After the overnight incubation, an additional 900 µL of 1:1 dichloromethane:methanol was added to the samples and incubated on a plate rotator for an additional hour before spinning samples at 2000 x g for 15 minutes to separate cell debris from the lipid extracts. The supernatant was transferred to a new 96-well plate for analysis by mass spectrometry as detailed below.

### Measurement of total sphingolipid levels using acid base hydrolysis

Blood samples and microbial cultures were broken down to their sphingolipid long chain base backbones using harsh acid and base treatment to obtain estimates of total sphingolipid levels using the methods outlined in (*38*). In brief, equal volume of hepatic portal vein blood (150 µL per experiment) or protein normalized microbial cultures were added to 500 µL of methanol supplemented with 4 µM of 1-deoxysphinganine-D3 (C_18_H_36_D_3_NO) (Avanti Polar Lipids) and vortexed on a plate vortexer for 1 hour. Samples were then spun down at 21130 x g in a microcentrifuge for 5 minutes to remove cell debris. Supernatants were transferred to polypropylene tubes and 75 µL of concentrated hydrochloric acid was added before incubating the samples overnight (16-20 hours) at 65°C. After overnight incubation, concentrated potassium chloride (10M) was added to samples and a lipid extraction using chloroform was used to extract hydrolyzed sphingolipids. The organic phase of the lipid extraction was dried under nitrogen gas and resuspended in 200 µL of 1:1 dichloromethane:methanol before adding C12 ceramide (d18:1/12:0) (Avanti Polar Lipids) internal standard to samples on a 96 well plate and analyzing sphingolipid levels by mass spectrometry.

### Lipidomic profiling

Sphingolipid abundance was measured by liquid chromatography-mass spectrometry (LC-MS). Specifically, 4 µl of lipid extract from each sample was injected into an Agilent 1200 HPLC (Agilent Poroshell 120 column) linked to an Agilent 6430 triple quadrupole mass spectrometer. Mobile phase A consisted of methanol/water/chloroform/formic acid (55:40:5:0.4 v/v); Mobile phase B consists of methanol/acetonitrile/chloroform/formic acid (48:48:4:0.4 v/v). After pre-equilibration for 6 sec, the gradient gradually increases to 60% mobile phase B and 100% mobile phase B that is held for 1.9 min. Flow rate was 0.6 mL/min. The duration of a run was 9.65 min. Ions were fragmented using electrospray ionization in positive mode and selective reaction monitoring (SRM) allowed for the detection of sphingolipid specific transitions. Peak calls and abundance calculations were done using MassHunter Workstation Software Version B.06.00 SP01/Build 6.0.388.1 (Agilent). Final concentrations of samples were calculated from a standard curve for each sphingolipid (Table S2). For Sa (d17:0) base metabolites, no standard curve was available and the response of the mass spectrometer normalized to the standard (Ceramide (d18:1/12:0)) was used for abundance calculations.

### Statistical Analyses

All data are represented as the mean ± SEM unless otherwise noted. Statistical tests are denoted in figure legends and were implemented using Prism 7.0 (GraphPad) or using the Tukey C, nlme, and multcomp packages implemented in R.

## Supplemental Figures and Figure Legends

**Figure S1.**
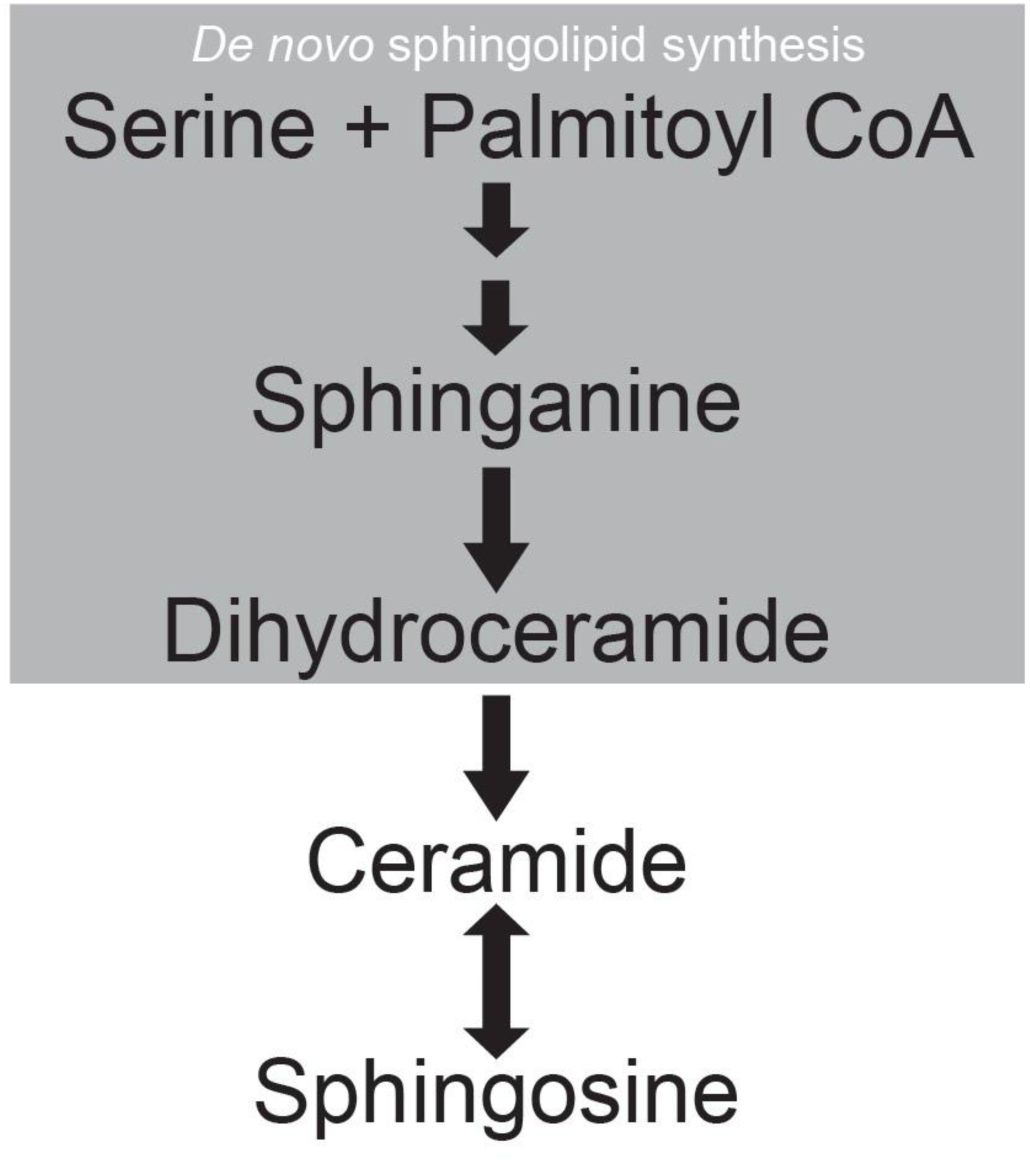
Sphingolipid synthesis pathway. SL synthesis pathway with the steps of *de novo* synthesis highlighted by the grey box.

**Figure S2.**
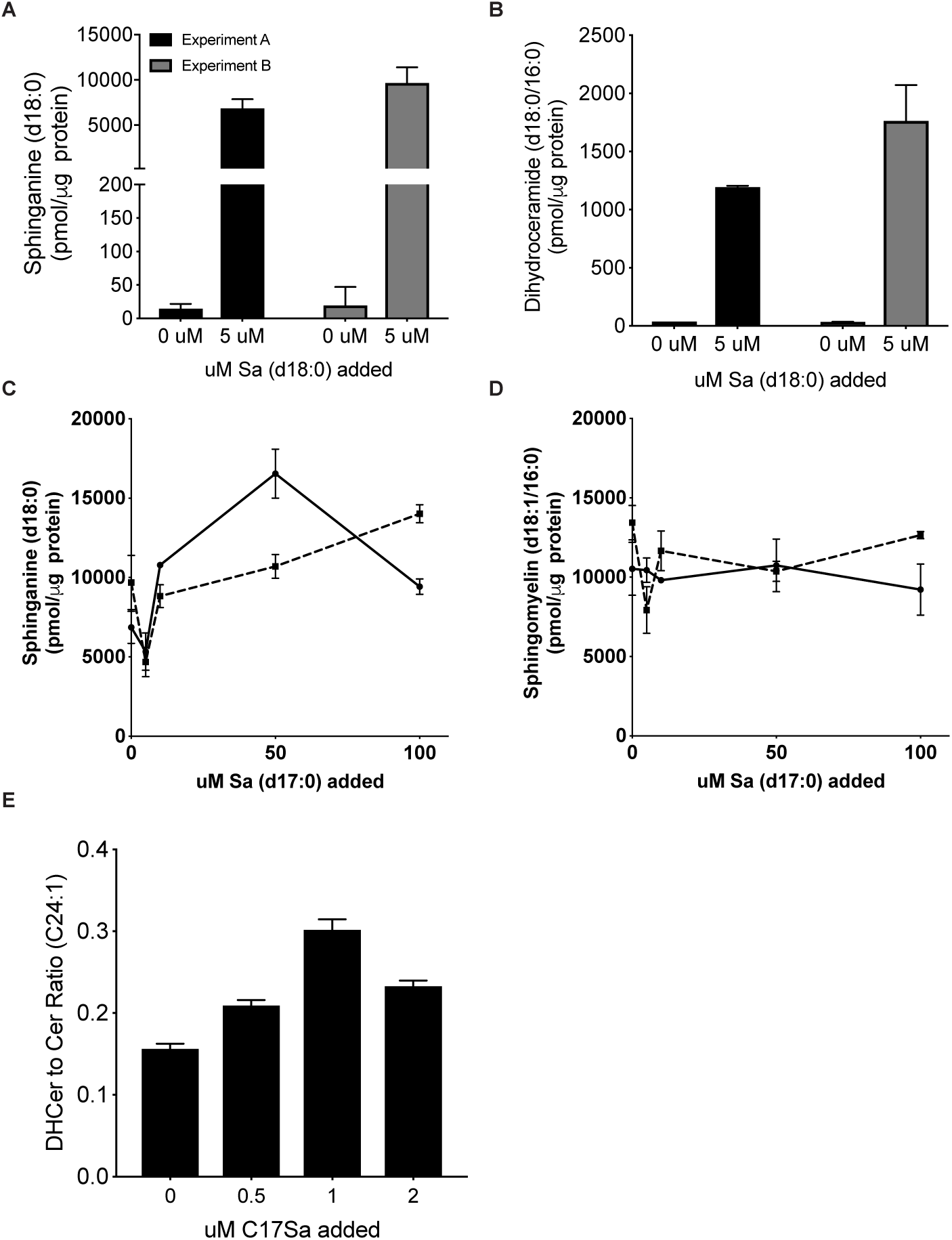
Sphinganine (d18:0) supplementation induces *de novo* sphingolipid synthesis in Caco-2 cells. (A-B) Levels of *de novo* sphingolipid synthesis pathway metabolites.(A) sphinganine (d18:0) and (B) dihydroceramide (d18:0/16:0) in Caco-2 cells with or without media supplementation with 5uM sphinganine (d18:0). (A-B) Means ± SEM of LC-MS measurements (n=2) are plotted for experiment replicates A and B. (C - D) Sphingolipid synthesis was induced in proliferating Caco-2 cells with addition of 5 uM Sa (d18:0) and then cells were dosed with increasing concentrations of Sa (d17:0) to monitor the ability of Sa (d17:0) to inhibit levels of C18-base sphingolipid species through the sphingolipid synthesis pathway. Cells were harvested 1 hour after addition of lipids and the two curves (dotted and straight lines) represent independent replicates of the same time course. Means ± SEM of LC-MS measurements (n=2) are plotted. (E) Amount of *de novo* sphingolipid synthesis of ceramide (d18:1/24:1) with increasing concentrations of sphinganine (d17:0) in Caco-2 cells. Means ± SEM of biological replicates (n=2) are plotted.

**Figure S3.**
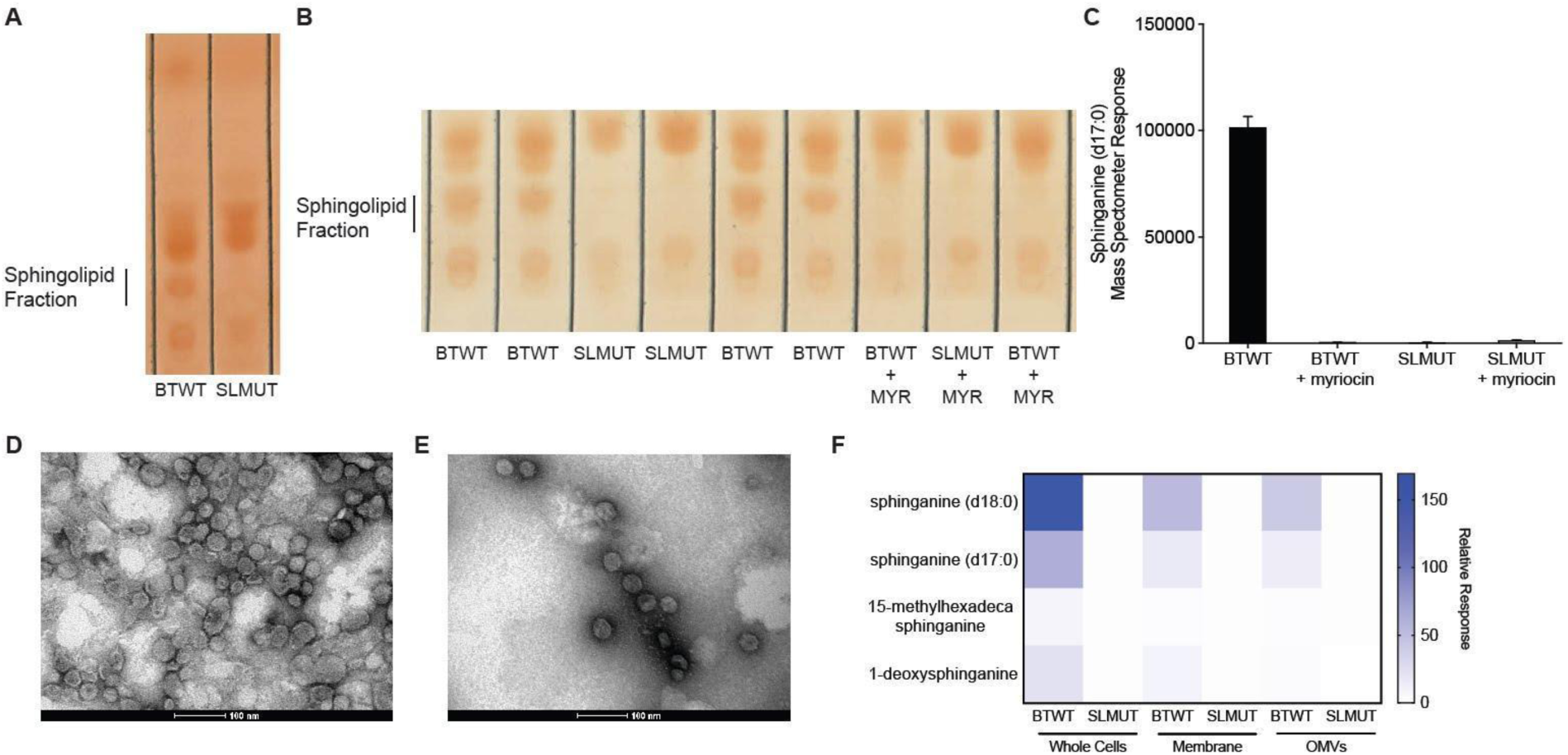
Genetic disruption of the putative serine palmitoyl transferase (SPT) gene by a transposon or drug treatment with the SPT inhibitor myriocin (myr) both inhibit sphingolipid synthesis in *Bacteroides thetaiotaomicron*. (A) Thin layer chromatography (TLC) of BTWT and SLMUT lipid extracts showing the SL fraction. (B) TLC of lipid extracts showing the SL fraction. Shown are BTWT (4 lanes), SLMUT (2 lanes), and each strain treated with myriocin (+ MYR; 2 lanes for BTWT and 1 lane for SLMUT), a chemical inhibitor of sphingolipid synthesis. (C) Mass spectrometry measurements of the major sphingolipid, sphinganine (d17:0), in BTWT and SLMUT strains treated or not treated with myriocin. Experiments were performed in duplicate and profiles represent measurements from one representative experiment. (D-E) Electron micrographs of OMVs fractionated from BTWT (D), and SLMUT (E) cultures. Purity of OMVs was confirmed by the presence of 20 – 250 nm sized particles from log phase cultures. Scale bar is 100 nm. (F) LC-MS based measurement of four sphingolipids identified in BTWT but absent in SLMUT for whole cells, membrane fractions, and isolated OMVs.

**Figure S4.**
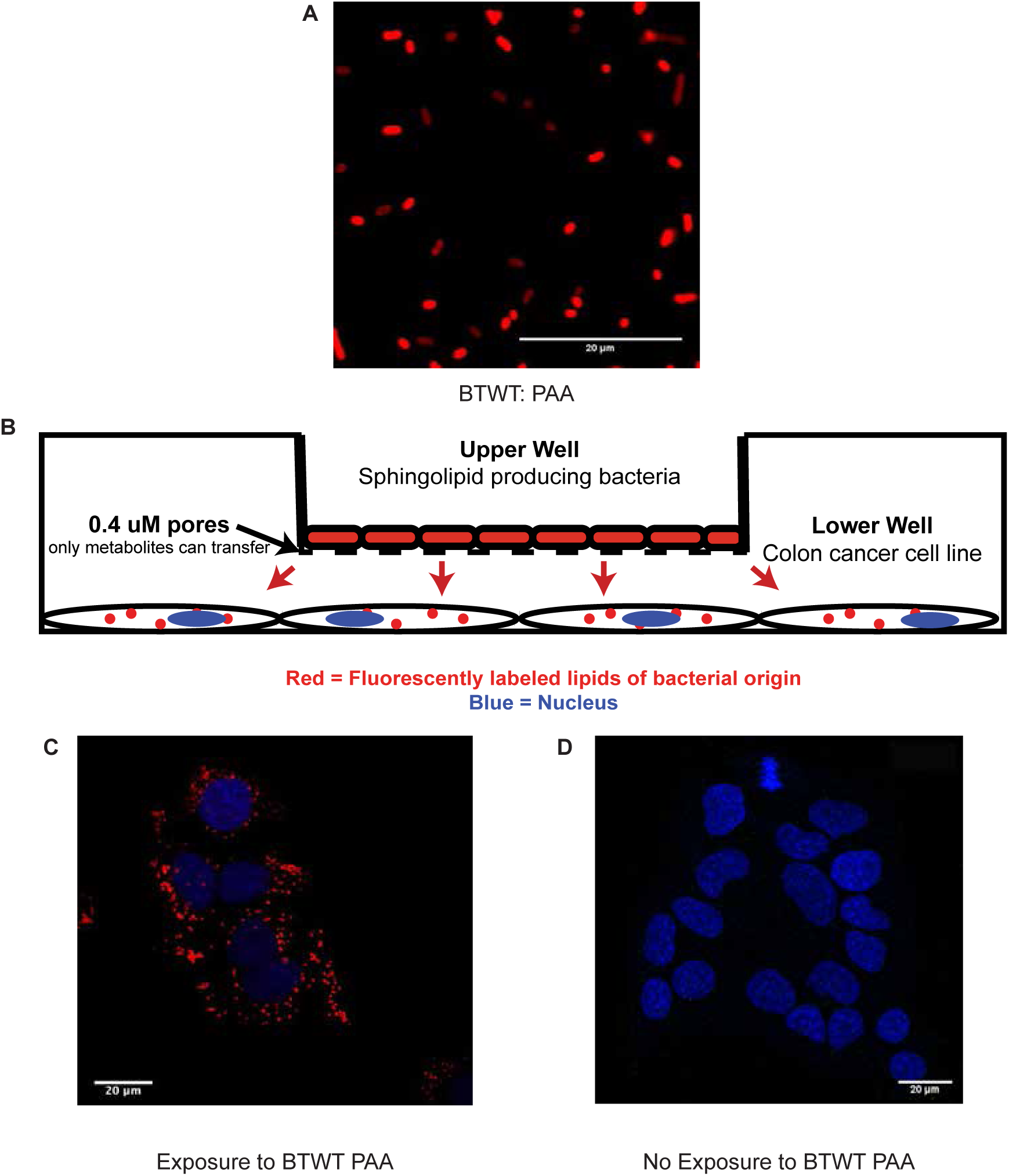
Lipids from BTWT transfer to Caco-2 cells in a transwell system. (A) Confocal image showing palmitic acid alkyne (PAA) or derivatives detected in BTWT cells grown with PAA (Red -Alexa Fluor 647; detection by click chemistry). Scale bar is 20 μm. (B) Cartoon diagram of the transwell coculture system, showing transfer of PAA or derivatives from BTWT-PAA in upper well to Caco-2 cells in lower well. (C) Confocal microscopy image of Caco-2 cells in the lower well after 6-hr exposure to BTWT-PAA. Red = alkyne tag; blue = DAPI. Scale bar is 20 μm. (D) Control for non-specific red fluorescence. Caco-2 cells unexposed to BTWT-PAA. Scale bar is 20 μm.

**Figure S5.**
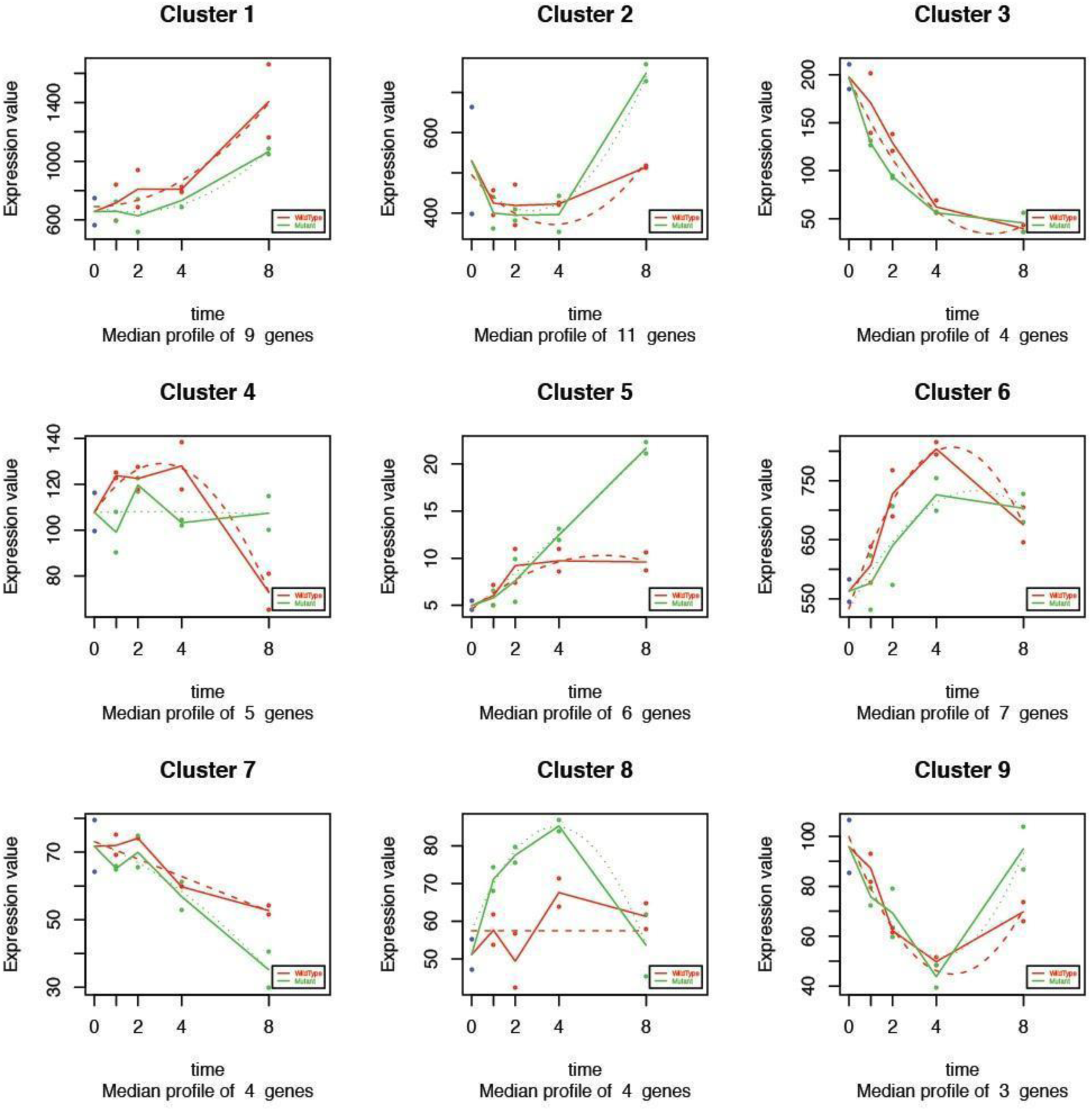
Gene expression changes in Caco2 cells cultured in transwell with BTWT or SLMUT over an 8-hour time course. Changes in gene expression were monitored over an 8-hour time period by RNA-seq. Time course measurements were made in duplicate. Clusters of genes with significantly different expression profiles in Caco-2 cells over the 8-hour time course between BTWT (WildType - red) and SLMUT (Mutant - green) incubated cells. The x-axis shows sequencing depth normalized read count values. Solid lines connect the average expression values within a condition over time and dotted lines are the regression fit. Categories of gene functions and gene names are included in Table 1.

**Figure S6.**
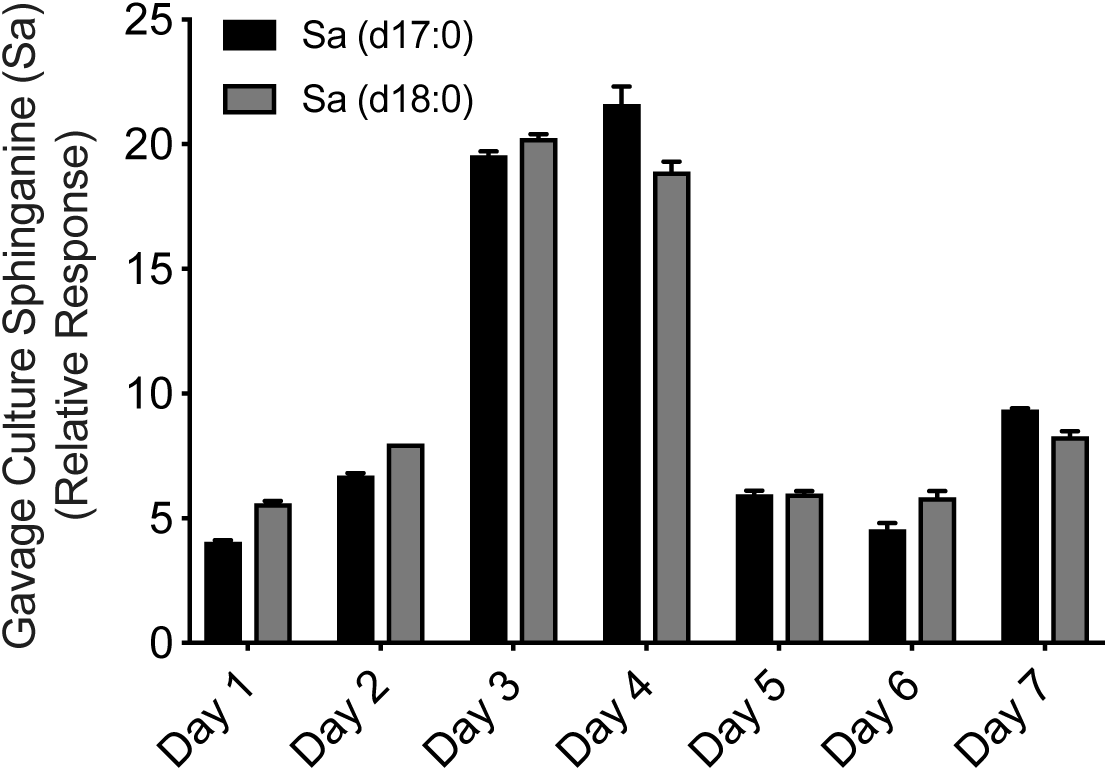
Sphinganine levels in BTWT cultures used to gavage germfree mice. Sphinganine (Sa (d17:0)) and sphinganine (Sa (d18:0)) - levels in BTWT cultures grown in minimal media as measured by LC-MS (relative response). Means ± SEM of LC-MS measurements (n=2) are plotted.

**Figure S7.**
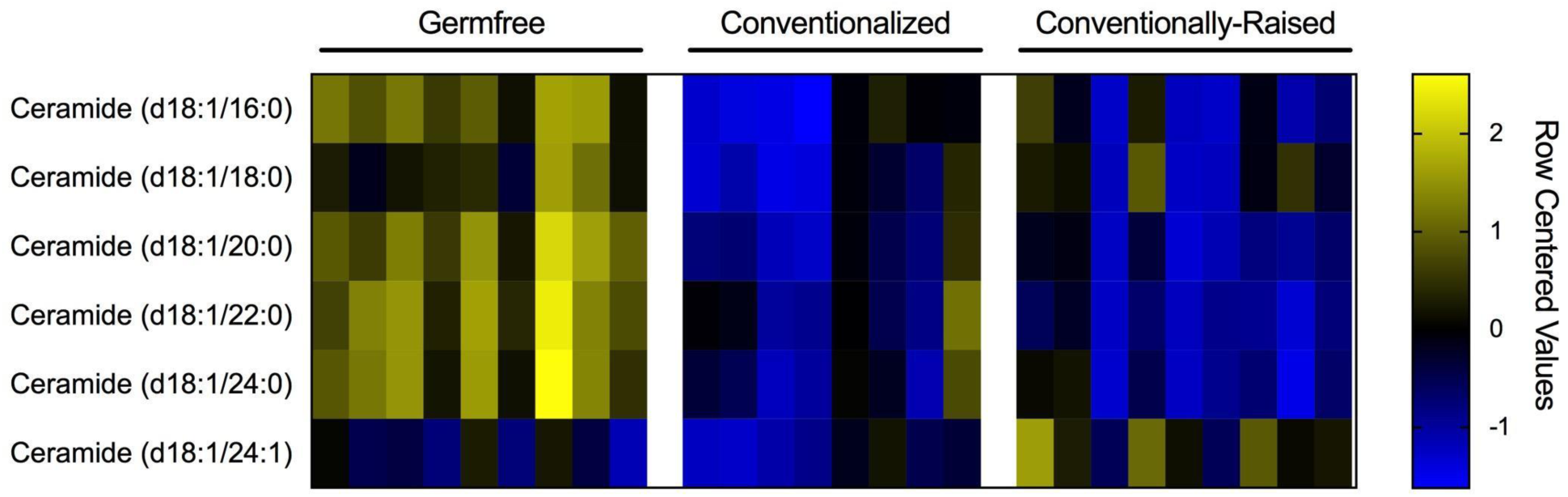
Hepatic sphingolipid levels in germfree, conventionalized, and conventionally-raised mice. Row centered values calculated from the LC-MS determined final concentration (pmol/ug protein) of hepatic sphingolipids in germfree mice (n=9, left), germfree mice one-week after introduction of a mouse microbiota (conventionalized, n=8, middle), and conventionally-raised mice (n=9, right).

**Figure S8.**
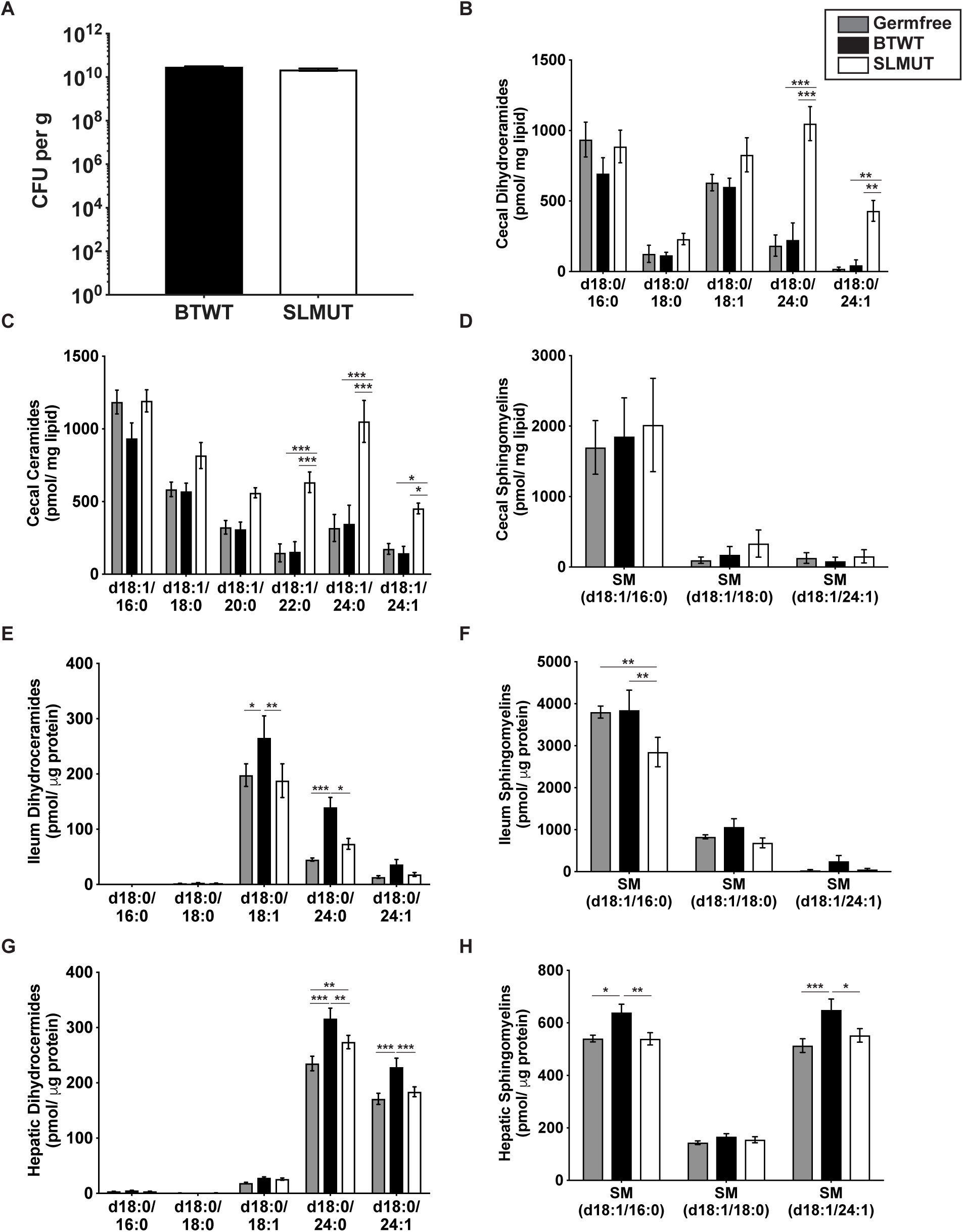
Colonization and sphingolipid levels of mice monoassociated with BTWT or SLMUT. (A) Colonization efficiency of BTWT and SLMUT strains gavaged into germfree mice in CFU/g as measured by serial dilution of cecal content. (B-H) Levels of SLs in different sample types obtained from: germfree mice (grey), germfree mice monoassociated with BTWT (black), and germfree mice monoassociated with SLMUT (white). (B) Cecal dihydroceramides; (C) cecal ceramides; (D) cecal sphingomyelins; (E) ileum dihydroceramides; (F) ileum sphingomyelins; (G) hepatic dihydroceramides; (H) hepatic sphingomyelins. Bar charts represent mean sphingolipid abundance ± SEM for 12 mice per condition (GF, SLMUT) and 11 mice per condition (BTWT), (two-way ANOVA, Tukey’s multiple comparison test, *p<0.05, **p<0.01, ***p<0.001).

**Figure S9.**
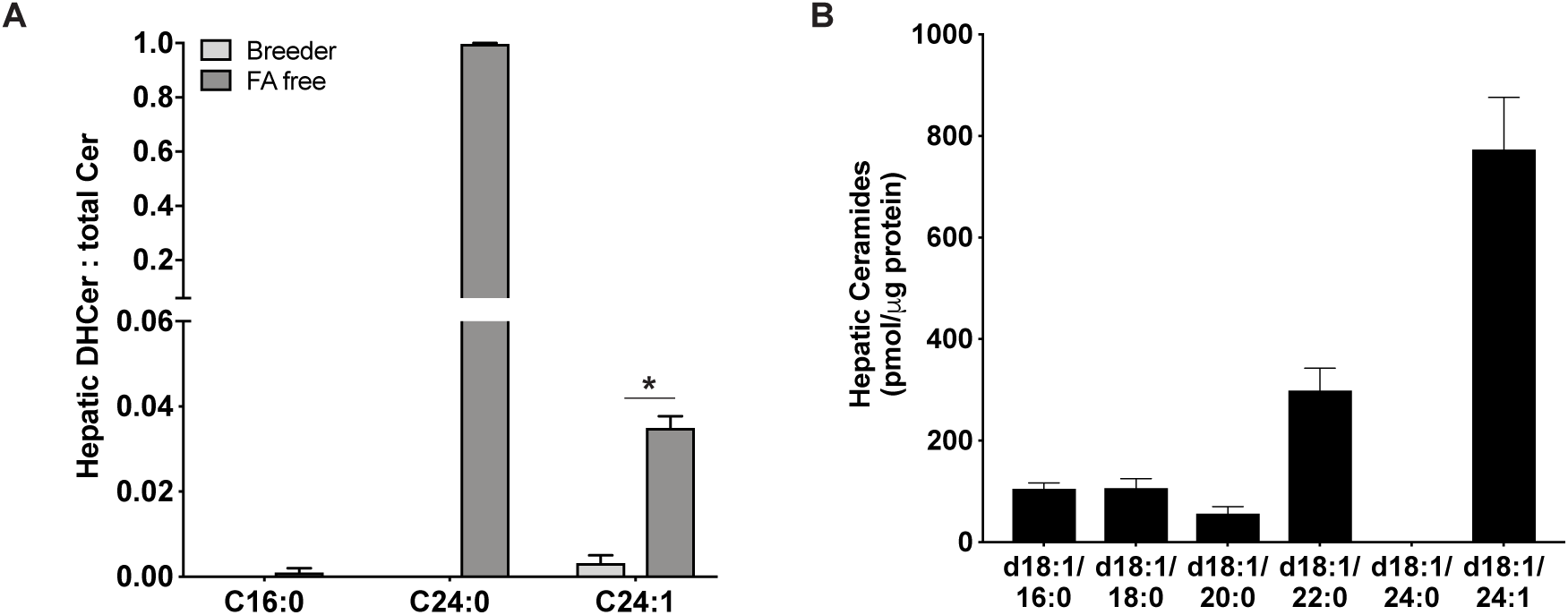
Hepatic *de novo* sphingolipid synthesis is induced and C_24:1_ is the major ceramide species detected in livers of mice on a fatty acid free diet. (A) Ratio of hepatic dihydroceramides to total ceramides(DHcer : total Cer) in livers of mice on a fatty acid free diet as compared to mice on a breeder diet. Values are mean ± SEM and n=3 mice per condition. Observations with significant differences in sphingolipid abundance are marked with stars (two-sided t-test, *=p <0.05). (B) Hepatic ceramide levels in mice fed a fatty acid free diet. Values are mean ± SEM and n=3 mice per condition.

**Figure S10.**
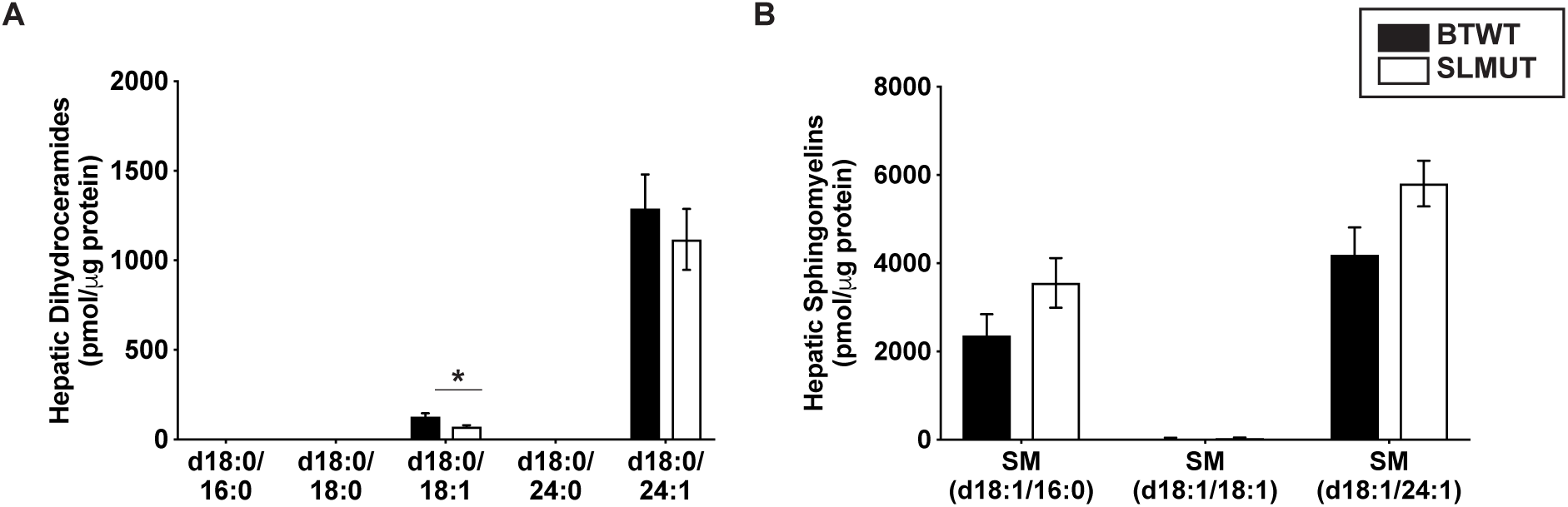
Sphingolipid levels in HFD-fed mice supplemented with BTWT or SLMUT. Hepatic dihydroceramides (A) and sphingomyelins (B) were measured at the conclusion of the IR experiment (A-B) Mean values ± SEM are plotted, n=10 per treatment, combined data from two replicate experiments with similar trends. (A-B) Observations with significant differences in SL abundance or glucose levels between treatments are marked with stars (t-test, *=p <0.05).

**Table S1.**
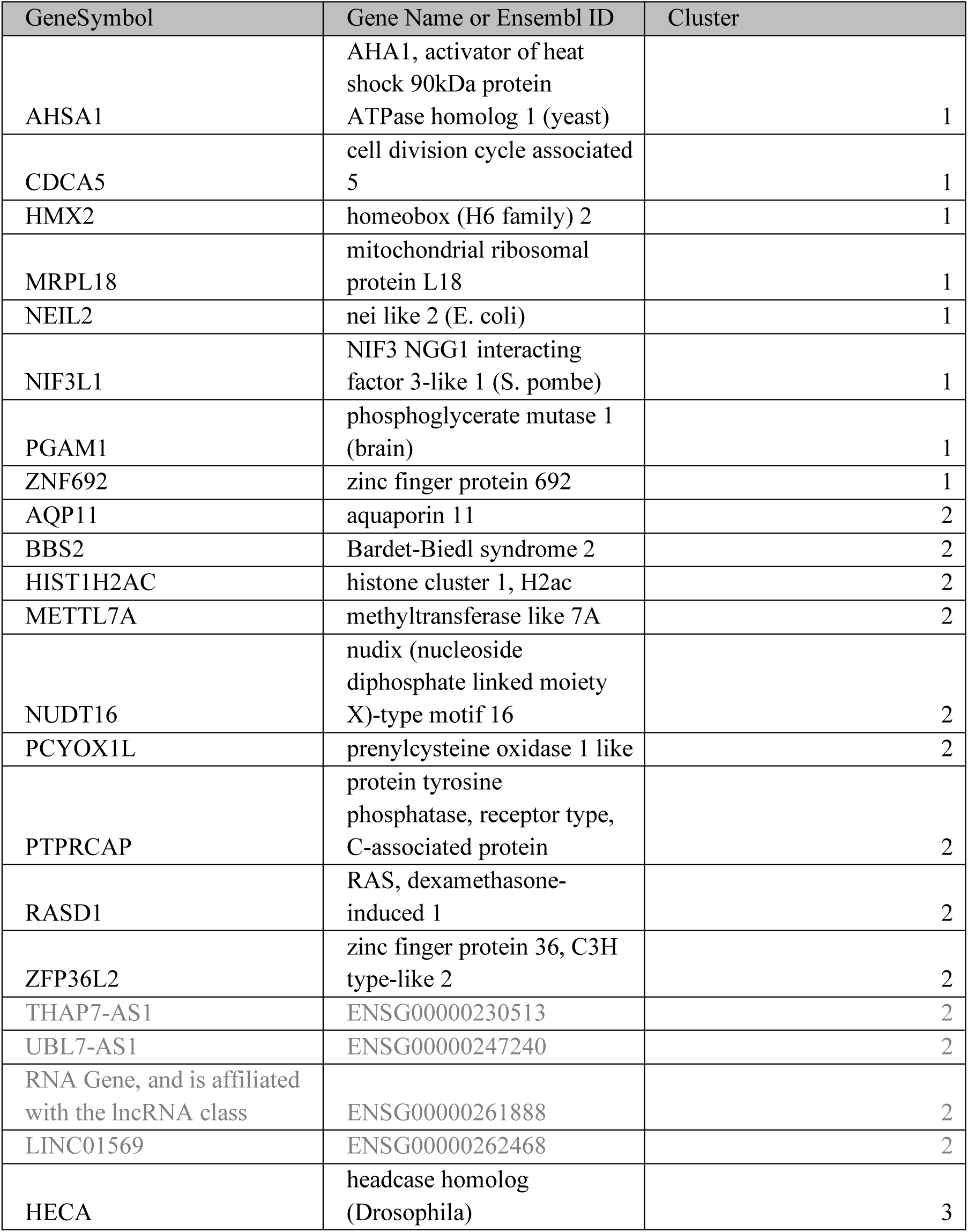
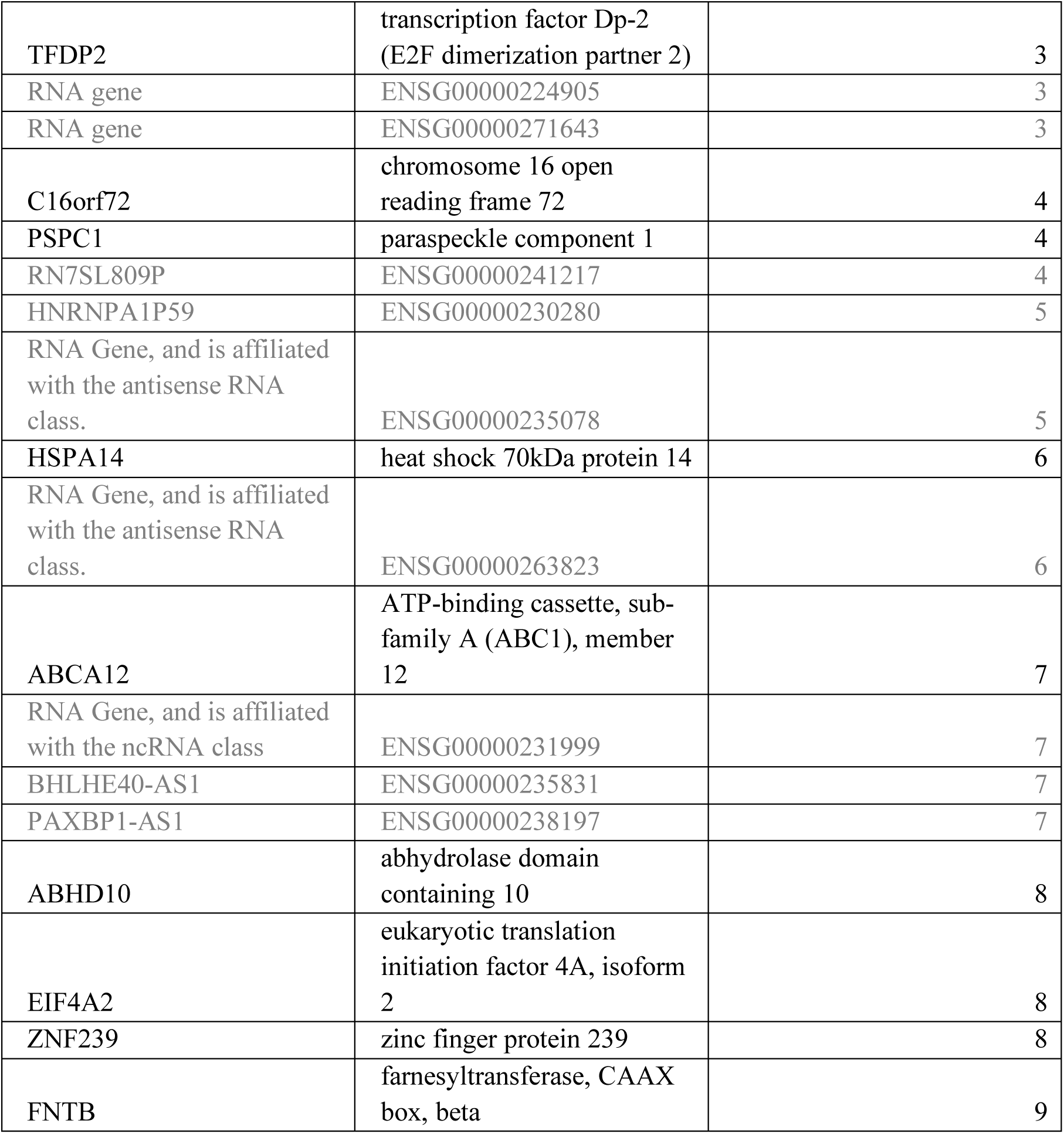
Genes with significant changes in expression profiles over an 8-hour time course in Caco-2 cells incubated with BTWT or SLMUT. Names and corresponding cluster identity of genes with significant differences in expression profiles between BTWT and SLMUT Caco-2 incubation conditions. Visualization of cluster expression profiles is shown in Fig. S5. Protein coding genes are in black while non-coding genes are in grey. Ten pseudo/unannotated genes are not listed.

**Table S2.** phingolipids measured by LC-MS in this study.

| **Sphingolipid Standard Name** | **Product Information** | **Mass Transition** |
| --- | --- | --- |
| Ceramide (d18:0/12:0) | https://avantilipids.com/product/860512 | Transition: 482.45 -> 464.5 |
| Ceramide (d18:1/16:0) | https://avantilipids.com/product/860516 | Transition: 538.5 -> 520.5 |
| Ceramide (d18:1/18:0) | https://avantilipids.com/product/860518 | Transition: 566.6 -> 264.3 |
| Ceramide (d18:1/20:0) | https://avantilipids.com/product/860520 | Transition: 594.6 -> 264.3 |
| Ceramide (d18:1/22:0) | https://avantilipids.com/product/860501 | Transition: 622.6 -> 264.3 |
| Ceramide (d18:1/24:0) | https://avantilipids.com/product/860524 | Transition: 650.7 -> 264.3 |
| Ceramide (d18:1/24:1) | https://avantilipids.com/product/860525 | Transition: 648.6 -> 264.3 |
| Dihydroceramide (d18:0/16:0) | https://avantilipids.com/product/860634 | Transition: 540.5 -> 522.5 |
| Dihydroceramide (d18:0/18:0) | https://avantilipids.com/product/860627 | Transition: 568.6 -> 550.6 |
| Dihydroceramide (d18:0/18:1) | https://avantilipids.com/product/860624 | Transition: 566.6 -> 548.5 |
| Dihydroceramide (d18:0/24:0) | https://avantilipids.com/product/860628 | Transition: 652.7 -> 634.6 |
| Dihydroceramide (d18:0/24:1) | https://avantilipids.com/product/860629 | Transition: 650.7 -> 632.6 |
| Sphingomyelin (d18:1/16:0) | https://avantilipids.com/product/860584 | Transition: 703.6 -> 184.1 |
| Sphingomyelin (d18:1/18:0) | https://avantilipids.com/product/860586 | Transition: 731.6 -> 184.1 |
| Sphingomyelin (d18:1/18:1) | https://avantilipids.com/product/860587 | Transition: 729.6 -> 184.1 |
| Sphingomyelin (d18:1/24:1) | https://avantilipids.com/product/860593 | Transition: 813.7 -> 184.1 |
| Sphinganine (d17:0) | https://avantilipids.com/product/860654 | Transition: 288.3 -> 270.3 |
| Sphinganine (d18:0) | https://avantilipids.com/product/860498 | Transition: 302.3 -> 284.3 |
| Sphingosine (d17:1) | https://avantilipids.com/product/860640 | Transition: 286.3 -> 268.3 |
| Sphingosine (d18:1) | https://avantilipids.com/product/860490 | Transition: 300.3 -> 282.3 |

**Table S3.**
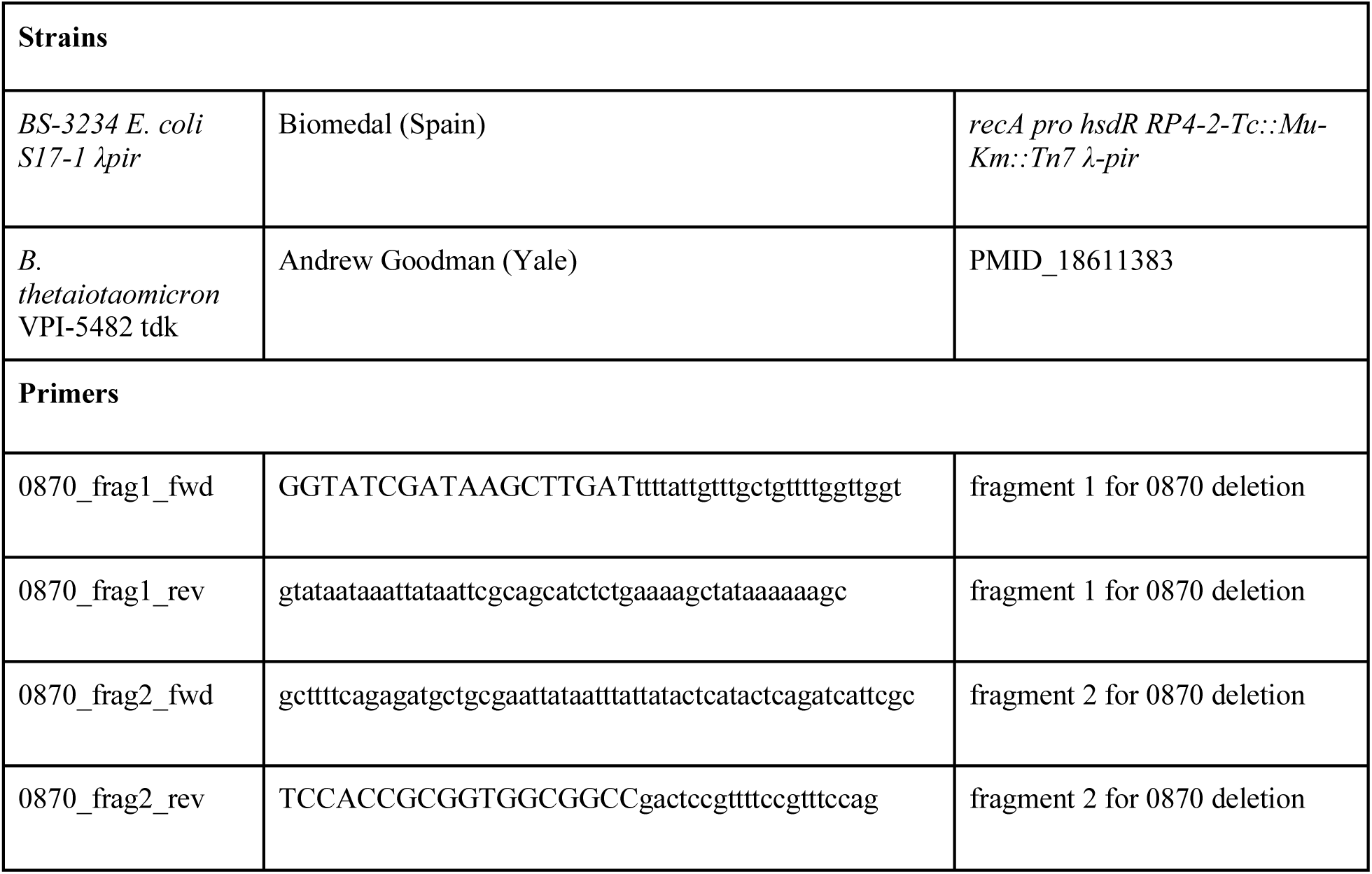
Bacterial strains and plasmids used in the generation of a BT0870 knockout strain.

Other supplementary material for this manuscript include:

Table S4 – RNA-seq read counts (excel)

Table S5 – Sphingolipid genes used in RNA-seq analysis (excel)

**Additional Data Table S4 (separate file)**

Excel file containing RNA-seq read counts

**Additional Data Table S5 (separate file)**

Excel file containing SL genes used in RNA-seq analysis

